# AutoMacq: an automatic pipeline to analyse macaque structural MRI data

**DOI:** 10.1101/2023.06.06.543846

**Authors:** Nathan Kindred, Susanna Carella, Martijn P van den Heuvel, Fabien Balezeau, Yujiang Wang, Colline Poirier

**Affiliations:** Biosciences Institute & Centre for Behaviour and Evolution, Faculty of Medical Sciences, Newcastle University, United Kingdom; Department of Complex Traits Genetics, Center for Neurogenomics and Cognitive Research, Vrije Universiteit Amsterdam, Amsterdam Neuroscience, Amsterdam 1081 HV, the Netherlands; Department of Child Psychiatry, Amsterdam University Medical Center, Location Vrije Universiteit Amsterdam, Amsterdam Neuroscience, Amsterdam 1081 HV, the Netherlands; Biosciences Institute, Faculty of Medical Sciences, Newcastle University, United Kingdom; CNNP Lab (www.cnnp-lab.com), Interdisciplinary Complex Systems Group, School of Computing, Newcastle University, United Kingdom

## Abstract

MRI scanning of rhesus macaques is a growing field due to their evolutionary proximity and similar neuroanatomy to humans. As such, there is a need for automatic macaque MRI processing pipelines. AutoMacq is a pipeline capable of processing rhesus macaque MRI data to produce both voxel-based and surface-based metrics. It involves minimal manual intervention and can be carried out without expert knowledge of macaque neuroanatomy. To test the quality of the pipeline, scans from 74 subjects across 8 different sites were processed. Results indicate that over 87% of tissue segmentations and surfaces were of satisfactory quality to not require additional manual correction. Hemispheric comparisons and analyses of scan-rescan data showed strong reliability of the volumetric and surface-based outputs. Finally, to illustrate potential applications of AutoMacq, the change in grey matter volume with ageing was investigated cross-sectionally using subjects aged 3-15 years (corresponding to adolescence until mid-adulthood). The analysis revealed a linear decrease in grey matter volume with age similar to what has been found in humans, reinforcing the value of rhesus macaques as a model of healthy human ageing.

## 1. Introduction

Automatic pipelines are the gold standard in MRI data processing as they avoid biases that could be introduced by manual interventions and allow for standardisation of data processing. Many pipelines have been created to automatically process and analyse human MRI data, with minimal manual intervention from the researcher (https://neuro-jena.github.io/cat/; https://surfer.nmr.mgh.harvard.edu/fswiki/FreeSurferAnalysisPipelineOverview; https://www.nipreps.org/smriprep/; https://www.fil.ion.ucl.ac.uk/spm/software/spm12/; Fischer *et al*. 2012; Glasser *et al*. 2013; Reuter *et al*. 2012). They can handle both cross-sectional and longitudinal datasets and allow for investigation of voxel-based morphometry (VBM) and/or surface-based morphometry (SBM).

Over the last couple of decades, MRI scanning of animal models has become a rapidly growing field (Öz, Tkáč and Ugurbill 2013). Non-human primates are of particular comparative and translational interest due to their relative evolutionary proximity to humans. In particular, their similarity to humans in terms of brain anatomy and cognitive abilities have made them crucial model animals in neuroscience research (Phillips *et al*. 2014; Roefsema and Treue 2014; Stonebarger *et al*. 2021). Additionally, Rhesus macaque are of particular value for ageing research due to their comparable life stages to humans, combined with their accelerated rate of ageing (3-4 times the rate of humans) which can allow for more efficient research (Mattison and Vaughan 2017).

The processing of macaque MRI data comes with unique issues, precluding the use of established human pipelines to process macaque MRI data (Milham et al., 2018; PRIMatE Data Exchange (PRIME-DE) Global Collaboration Workshop and Consortium 2020). Though similar in shape and organisation to the human brain, the macaque brain is around 12-16 times smaller than the human brain (in terms of volume), with specific areas accounting for different proportions of the macaque brain compared to the human brain (Croxson *et al*. 2018). Other differences include differences in the amount of tissue surrounding the brain as well as in tissue contrast, making it more difficult to extract and segment the brain in MRI scans, compared to human scans. Non-standardized surface coil arrangements are common when scanning macaques, and often result in variations in coil coverage and image intensity. Differences between sites in terms of equipment and protocols are also common and result in data across sites that varies greatly in terms of quality and scan parameters (Milham *et al*. 2018). Furthermore, motion artefacts can be an additional issue when scanning awake macaques, with the only current way to minimise these artefacts being through training and/or head fixation (PRIMatE Data Exchange (PRIME-DE) Global Collaboration Workshop and Consortium 2020).

Custom processing pipelines tailored for rhesus macaque MRI data are clearly required, and over the last few years such pipelines have been developed (Balbastre *et al*. 2017; Garcia-Saldivar et al. 2021; Lepage et al. 2021). These pipelines require manual correction of tissue segmentation and surfaces, which relies on expert knowledge of macaque neuroanatomy. Additionally, the currently available macaque processing pipelines implement surface-based morphometry only, with no clear option to carry out voxel-based morphometry, despite the complementarity of the two approaches (Goto *et al*. 2022).

This study therefore aimed to design a processing pipeline for rhesus macaque MRI data that can produce accurate tissue segmentations and surfaces without manual intervention and can produce both voxel-based and surface-based metrics.

## 2. Materials and Methods

### 2.1. Datasets

Scans from publicly available datasets, as well as those acquired locally or privately shared with the authors, were selected to test the pipeline. Data had been acquired on various scanners with various parameters and coil arrangements. An initial visual quality control of the scans was carried out prior to AutoMacq processing and the following section describes the datasets that were retained (for excluded scans and justification, see Suppl. Fig 1).

#### Cross-sectional datasets

The AutoMacq pipeline was tested using cross-sectional T1 scans from 8 different sites (N= 74 subjects, see Table 1; detailed scan parameters for each site are provided in suppl. Table 1). One of these datasets was collected at Newcastle University, as part of an ongoing longitudinal project. Three other datasets came from Deutsches Primatenzentrum (DPZ), Germany, the National Institute on Drug Abuse (NIDA), USA, and the University of Oxford, UK. The remaining five cross-sectional datasets were from the primate data exchange (PRIME-DE) (Milham *et al*. 2018): Mount Sinai School of Medicine (MSP and MSS), USA, Stem Cell and Brain Research Institute (SBRI), France, University of California Davis (UCD), USA and University of Western Ontario (UWO), Canada. For some subjects, T2 data were available alongside the T1 scan (see Table 1). All the scans were acquired with a scanner strength of at least 3T and comprised a combination of scans acquired in anesthetized and awake animals.

**Table 1:**
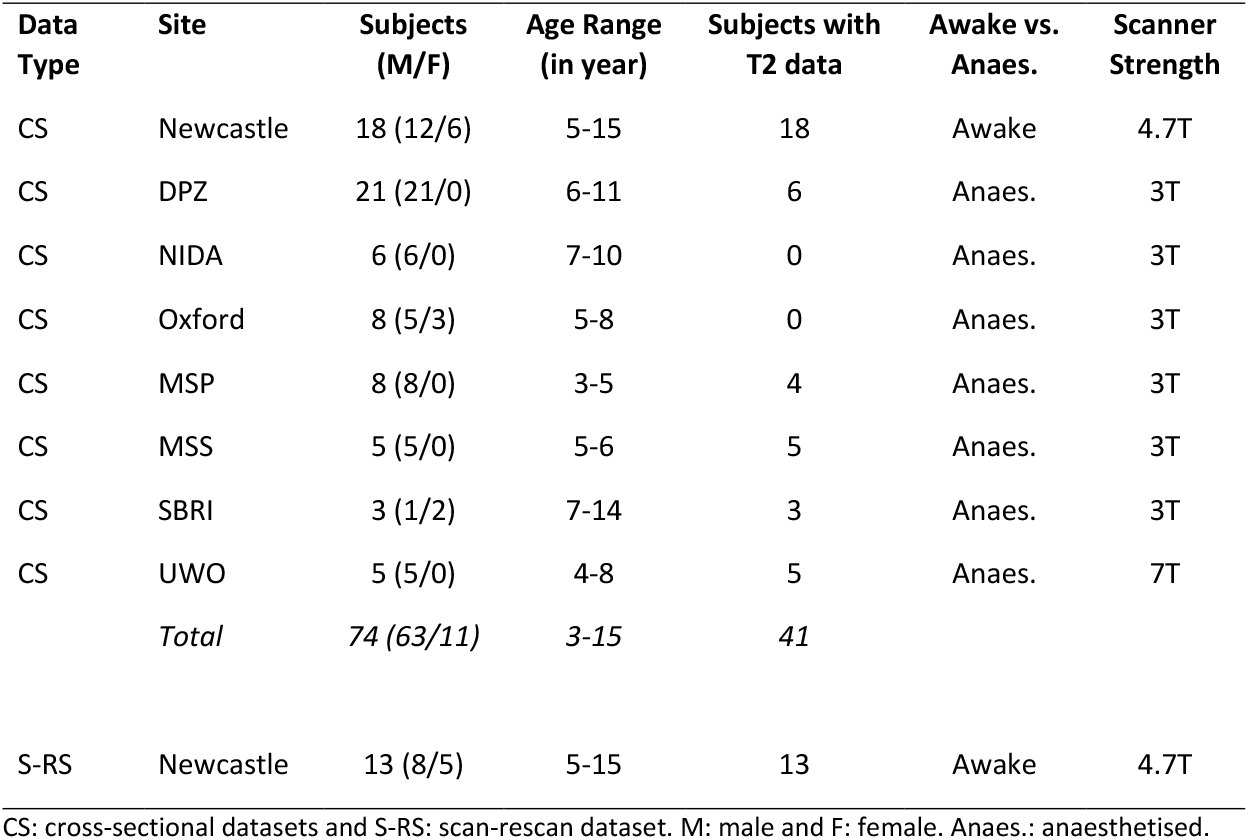
Description of included datasets after the initial quality control check.

#### Scan-Rescan datasets

For a subselection of data two scans per subject, acquired within one week, were available (N = 13). All data were acquired in awake animals, at Newcastle University. There was some overlap in the scans included in the cross-sectional and scan-rescan datasets from Newcastle University.

### 2.2. Ethics

Original data were acquired in accordance with the relevant legislation in each country and approved by an ethics committee. The re-use of the data was approved by Newcastle University Animal Welfare Ethical Review Board (reference number 1021).

### 2.3. Cross-sectional AutoMacq Pipeline

AutoMacq is optimised for the input of both T1 and T2 images, but can process T1 data alone, and utilises freely available software packages (SPM-https://www.fil.ion.ucl.ac.uk/spm/software/spm12/; FreeSurfer-Fischl 2012; ANTs-Avants *et al*. 2009; FSL-Jenkinson *et al*. 2012; Connectome Workbench-Marcus *et al*. 2011). The pipeline can use any macaque template. For this study, the population-average 112RM-SL template and its prior maps (McLaren *et al*. 2009) were chosen for testing the pipeline. This template is aligned to the Saleem-Logothetis atlas (Saleem and Logothetis 2012) that provides both high-resolution MRI scans and histological sections to delineate the anatomy of the macaque brain. An Ear Bar Zero (EBZ) coordinate system is employed, meaning that the origin is set to the midpoint of the interaural line (Saleem and Logothetis 2012).

The AutoMacq pipeline is outlined in figure 1, and each step to process cross-sectional data is described in detail below (with step numbers referring to those in Fig. 1). AutoMacq can also process longitudinal structural MRI data from rhesus macaques, and this is discussed in the supplementary materials (see suppl. text, suppl. Table 2 and suppl. Figure 2). A detailed walkthrough, scripts and SPM batches for AutoMacq are available at: https://github.com/Nsk97/AutoMacq.git

**Figure 1:**
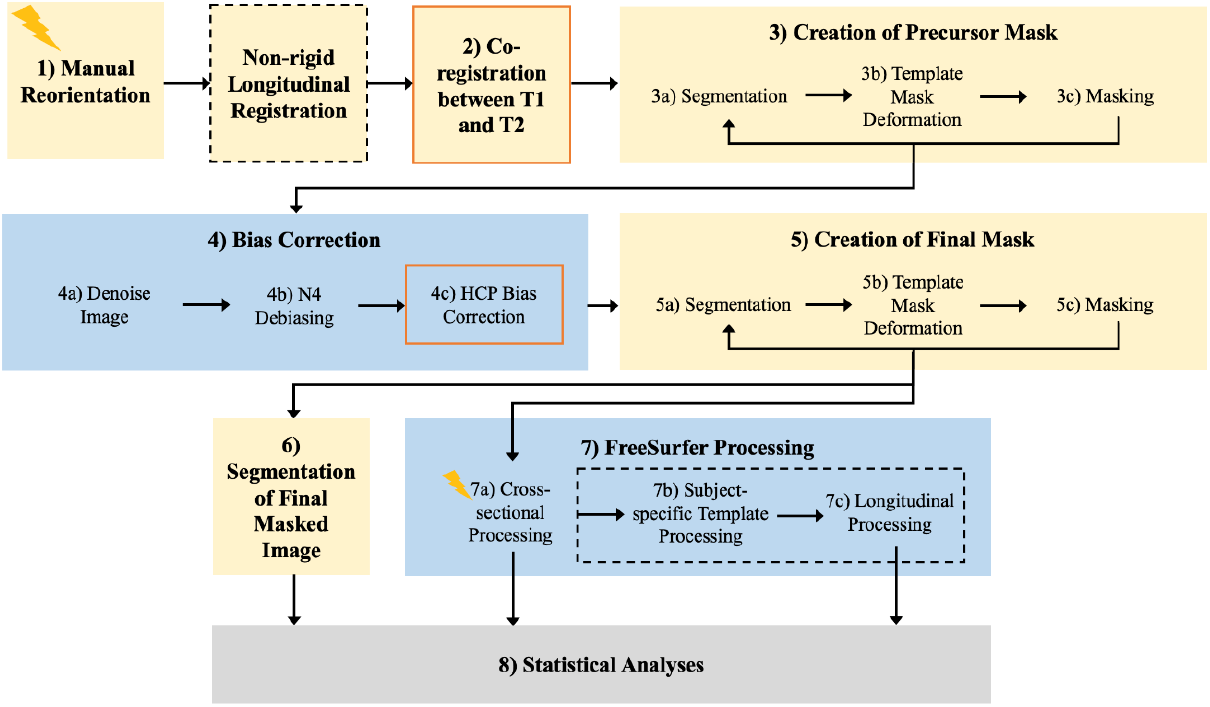
The AutoMacq pipeline. Yellow boxes represent steps carried out using SPM, blue boxes those carried out in a Linux environment (with ANTs, connectome workbench, FSL and FreeSurfer installed). An orange outline represents a step that is skipped if T2 data are not available, and a dashed black outline a step that would only be carried out for longitudinal data. Lightning bolts indicate a manual intervention.

#### 2.3.1. Cropping and Manual Reorientation (step 1)

Prior to processing through the AutoMacq pipeline, scans with large fields of view are cropped in FreeSurfer. This cropping minimises empty space and tissue outside of the skull. Cropping the field of view allows for more accurate masking later in the pipeline.

Following this, every T1 and, if available, T2 scan is manually reoriented in SPM. This involves rotating the scans and setting the origin to match the orientation and origin of the atlas. This manual reorientation step is simple and does not require any knowledge of macaque neuroanatomy.

#### 2.3.2. Co-registration between T1 and T2 (step 2)

The next step of AutoMacq consists of co-registering the reoriented T1 and T2 scans to ensure that their orientations precisely match. This step is done using the SPM intra-subject co-registration routine using a rigid-body model and image reslicing (moving the T2 scan to align it with the T1 scan). This step is skipped in the absence of T2 scans.

#### 2.3.3. Brain extraction

To obtain accurate tissue segmentation of macaque data, it is helpful to first mask out non-brain tissues, a process called brain extraction or skull stripping. This is done in 3 steps: (1) the creation of a precursor mask in SPM; (2) bias correction carried out using ANTs, connectome workbench and FSL; and (3) the creation of the final mask in SPM.

##### Creation of Precursor Mask and initial brain extraction (step 3)

The precursor mask is an approximate, subject-specific mask that can be utilised for bias correction. A precursor mask in the native space is obtained by first creating a mask of the template (by binarising the sum of the grey matter (GM), white matter (WM) and cerebrospinal fluid (CSF) prior maps), and then deforming the template mask to match the subject-specific scan(s). This deformation between the template space and the native space is calculated using the SPM segmentation routine. This routine combines tissue classification of the subject-specific scans (combining information from T1 and T2 scans to improve the segmentation accuracy), correction of intensity non-uniformity (bias correction) and non-rigid co-registration to the template. The segmentation relies on 4 classes of tissue probability maps: GM, WM, CSF, and non-brain tissues (adding the non-brain tissue class was found to improve the quality of the segmentation). The T1 scan (and coregistered T2 scan, if available) is then masked using this approximate precursor mask.

This process (segmentation, mask deformation and masking) can be repeated several times if necessary to further increase the quality of the subject-specific mask, as long as no brain tissues are masked out. Macaque data differs from human in that images are often acquired using non-standardized arrangements of surface coils (Milham et al., 2018), generating increased variation in image intensity. Using the bias correction developed for human data in SPM, no parameter was found to be good enough to produce accurate subject-specific masks. To increase the quality of the mask, the precursor mask was created using little debiasing (heavy regularisation) in SPM and the main debiasing done outside SPM (please note that the precursor mask needs to be resliced to be used by other software).

##### Bias Correction (step 4)

For the AutoMacq pipeline, the ANTs functions DenoiseImage and N4BiasFieldCorrection are utilised. DenoiseImage removes noise from the scans using a spatially adaptive filter, and N4 debiasing is a variant of non-parametric, non-uniform, normalization (N3) debiasing (Tustison *et al*. 2010). DenoiseImage needs to be applied to the unmasked T1, whereas N4BiasFieldCorrection utilises an unmasked T1 image and the precursor mask as inputs.

To further minimise bias in the images, the program connectome workbench along with the bias correction script from the Human Connectome Pipeline (HCP) can be utilised for subjects with both T1 and T2 scans. This script uses the square root of T1w*T2w in order to correct the bias field, and improvements can be seen when this is used alongside other debiasing steps. The HCP script requires both unmasked and masked T1 images as inputs; the masked T1 is obtained by applying the function fslmaths (from FSL) to the N4 debiasing output.

##### Creation of Final Mask and Brain extraction (step 5)

A final mask is then created using the same approach described in step 3 but using the debiased scan(s) as input(s) of the segmentation, and a final masked T1 (and T2 if available) is produced which excludes non-brain tissues.

#### 2.3.4. Tissue segmentation (step 6)

A final segmentation of the masked, debiased T1 scan is then performed in SPM. The output files from this segmentation can then be used to calculate tissues volumes as well as the local amount (or density) of grey matter in each voxel. These metrics can then be analysed statistically for voxel-based morphometry studies.

#### 2.3.5. Surface-based Cross-Sectional Processing (step 7)

Surface-based processing in AutoMacq utilises custom analysis scripts that adapt the FreeSurfer standard processing stream for human MRI data. For cross-sectional processing, the FreeSurfer stream consists of 3 major stages: autorecon1, autorecon2 and autorecon3. For cross-sectional processing in AutoMacq, modifications are made to the autorecon1 and autorecon2 stages.

##### Autorecon1

Autorecon1 begins with computation of the affine transformation from the final masked T1 obtained in step 5 to the MNI305 atlas. This is required as atlas coordinates of different brain areas are needed for several downstream functions. The MNI305 is a human brain atlas, so this automatic computation tends to be extremely inaccurate for macaque data, and there is no simple way to substitute a macaque atlas for the MNI305 atlas. However, macaque MRI data can be successfully processed through FreeSurfer by manual correcting the atlas registration. This is done by matching the size and orientation of the masked T1 to the MNI305. Therefore, this manual step does not rely on any knowledge of macaque neuroanatomy and can be carried out quickly and easily by a non-expert. The rest of the autorecon1 stage includes correction of any remaining non-uniformity or fluctuations in intensity (unchanged FreeSurfer standard step). The final step of skull stripping is skipped since images have already been brain extracted in SPM.

##### Autorecon2

The autorecon2 stage begins with the segmentation of subcortical structures and the computation of their respective volumes. In AutoMacq, the standard stream is adapted to use the manually corrected atlas registration to initialise the subcortical segmentation. Next, WM is segmented to give a WM volume image (cerebellum and brainstem are excluded), which is then used to create the surface encoding the boundary of WM and GM in each hemisphere. These left and right WM surfaces are used as a starting point to generate surfaces encoding the boundary of GM and CSF in each hemisphere (‘pial surfaces’). However, this processing in FreeSurfer alone does not always produce accurate surfaces. Instead of manually correcting the white matter surfaces (WM edits), the WM segmentation file produced by SPM (step 6) can be used to re-run the cortical surface generation. To be recognised correctly in FreeSurfer, the WM segment image is first binarized in SPM (threshold of 0.2). A custom script was written to recognise this binarized WM volume as WM edits. As the last step of autorecon2, a binary mask of the cortical ribbon is then created.

##### Autorecon3

The autorecon3 stage carries out the co-registration of the GM and WM surfaces to the spherical atlas (spherical morph), in order to label brain regions for cortical parcellation. The entirety of autorecon3 can be ran unchanged from the standard FreeSurfer stream to acquire global brain metrics, but it is also possible to easily replace the human atlas with a macaque parcellation schema in order to acquire macaque parcellations.

### 2.4. Statistics

Whole brain measures of GM, WM and CSF were extracted using SPM for every subject. Hemispheric measures of the same metrics were also extracted in SPM, using a hemisphere mask created through manual editing of the Saleem-Logothetis atlas mask. Whole brain and hemispheric measures of grey matter and white matter surface area, as well as cortical thickness were taken from FreeSurfer for every subject. Intraclass correlation coefficient (ICC) was calculated in R, using an absolute-agreement, single-measurement, two-way mixed-effects model, for scan-rescan comparisons. Pearson correlation analyses were carried out in R (https://www.r-project.org/) for hemisphere comparisons. This was used rather than ICC for the hemisphere comparisons as ICC accounts for systematic offsets, which could occur biologically between hemispheres.

The change in GMV with ageing was also investigated in R. Tests for normality and heteroscedasticity were first carried out. A non-linear fit was found to not significantly improve the percentage of variance explained by the model, so a linear model was fitted. Site/scanner was included as a random effect in the model, and total intracranial volume (TIV) was controlled for (fixed effect) in order to account for differences in head size (Whitwell *et al*. 2001).

## 3. Results

### 3.1. VBM outputs

All of the cross-sectional scans processed through AutoMacq (N=74) produced an accurate brain mask for skull striping (suppl. figure 3). Figure 2 illustrates a representative example of tissue volume outputs (grey matter and white matter volumes) obtained as the result of SPM segmentation. 95.9% of scans (N=71/74) produced tissue volumes of comparable quality to those in figure 2. The remaining scans (4.1%, 3/74) resulted in some errors in the segmentation of grey and white matter (suppl. figure 4).

**Figure 2:**
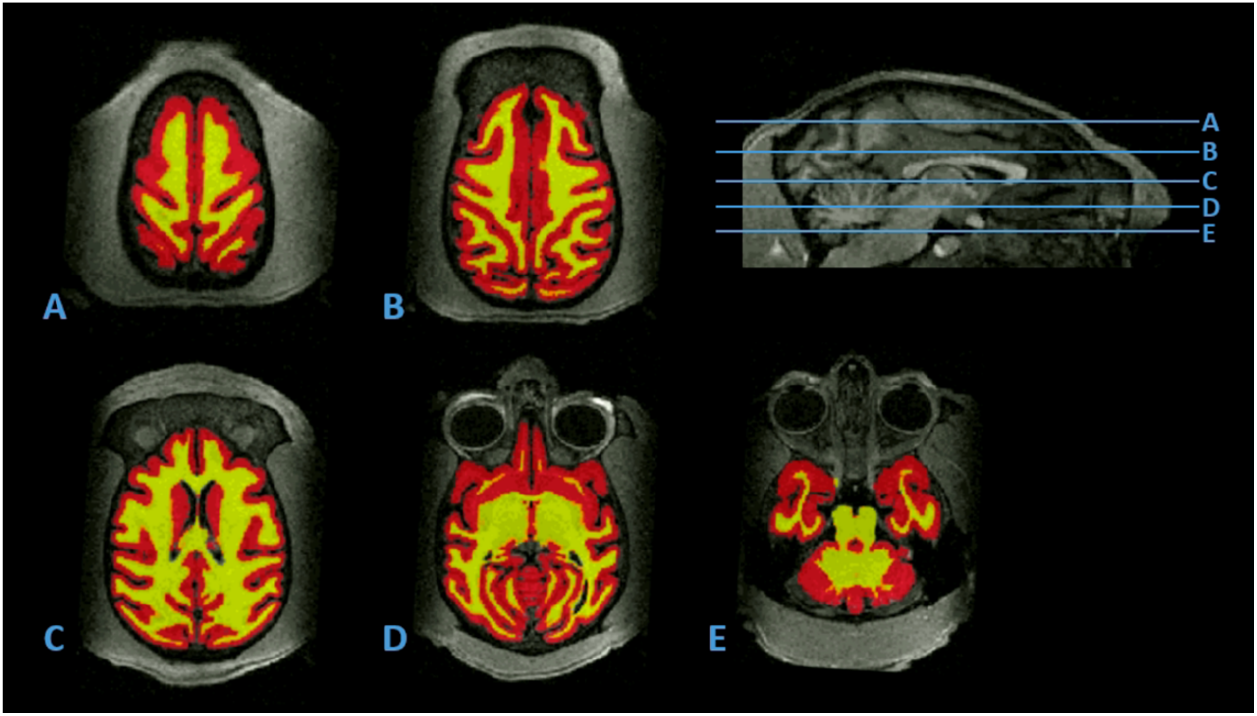
Representative tissue volumes produced by AutoMacq. Horizontal slices of a representative example of GMV (shown in red) and WMV (shown in yellow), produced using the cross-sectional AutoMacq pipeline, displayed on the corresponding T1 scan, and presented alongside a midsagittal image showing where each slice is taken from.

### 3.2. SBM output

Figure 4 shows a representative example of FreeSurfer surfaces output from the cross-sectional AutoMacq pipeline, for the same subject for which tissue volume outputs were displayed in figure 2. 87.8% of scans (65/74) processed through AutoMacq resulted in surfaces comparable to those shown in figure 4. This was after the WM segmentation file from SPM was used in place of WM edits in FreeSurfer for 60/74 (81.1%) subjects (the other 14 subjects produced good quality surfaces without this step). This improvement of the surface accuracy with this step is illustrated in figure 5. However, even after the incorporation of the WM segmentation file from SPM, 9/74 (12.2%) subjects still showed some errors in their surfaces (see details in suppl. figure 5).

**Figure 4:**
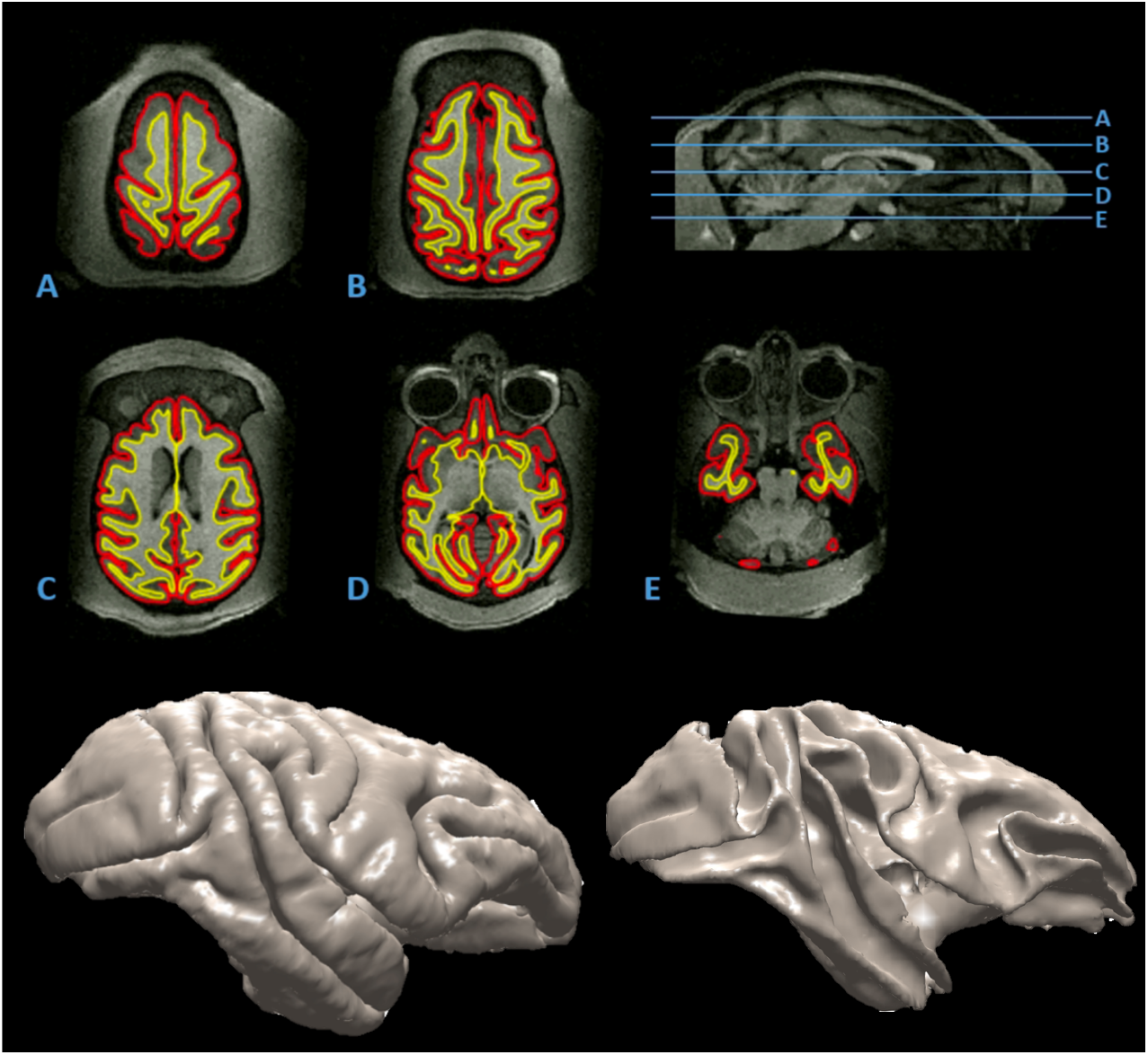
Representative surfaces produced by AutoMacq and 3D models of those surfaces. Horizontal slices of a representative example of pial (shown in red) and white matter (shown in yellow) surfaces, produced using the cross-sectional AutoMacq pipeline. A midsagittal image showing where in the brain each slice is taken from, and 3D models of the surfaces are also presented. The cerebellum and brainstem are excluded during FreeSurfer processing.

**Figure 5:**
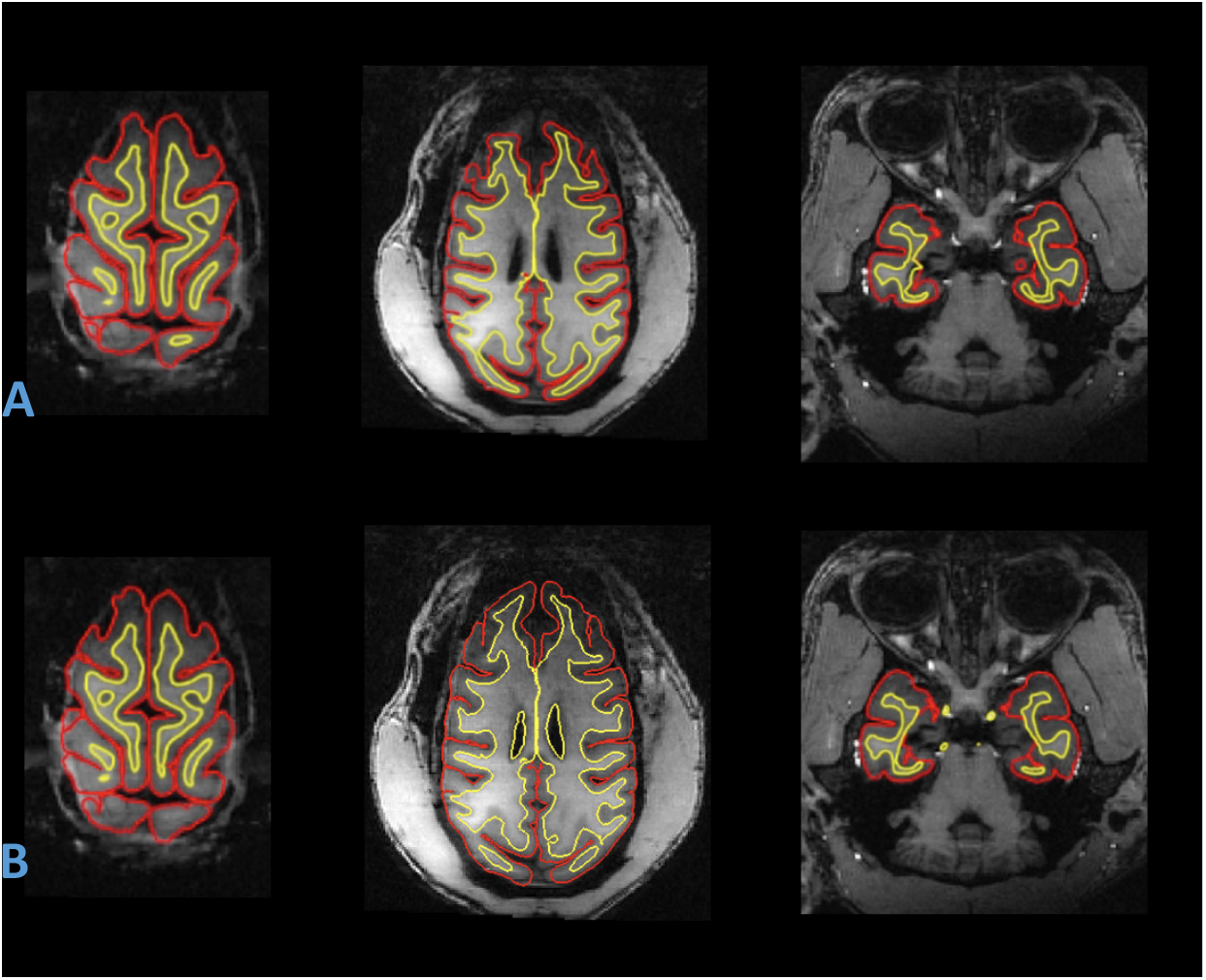
Comparison of surfaces produced with AutoMacq, without and with SPM WM. Horizontal slices of examples of pial (shown in red) and white matter (shown in yellow) surfaces, produced for the same subject using the cross-sectional AutoMacq pipeline, without WM from SPM (A) and with WM from SPM (B).

### 3.3. Hemisphere Comparison

To assess the reliability of AutoMacq, various volume-based and surface-based metrics were compared between hemispheres. Considering that hemispheric differences from biological origin are minimal, this analysis allows for quantification of errors mainly due to AutoMacq processing. All of the subjects processed through AutoMacq (including those with errors in their outputs) were included in this analysis.

Results indicate strong correlation between hemispheres for all metrics tested (R values between 0.9 and 0.98, Fig. 6).

**Figure 6:**
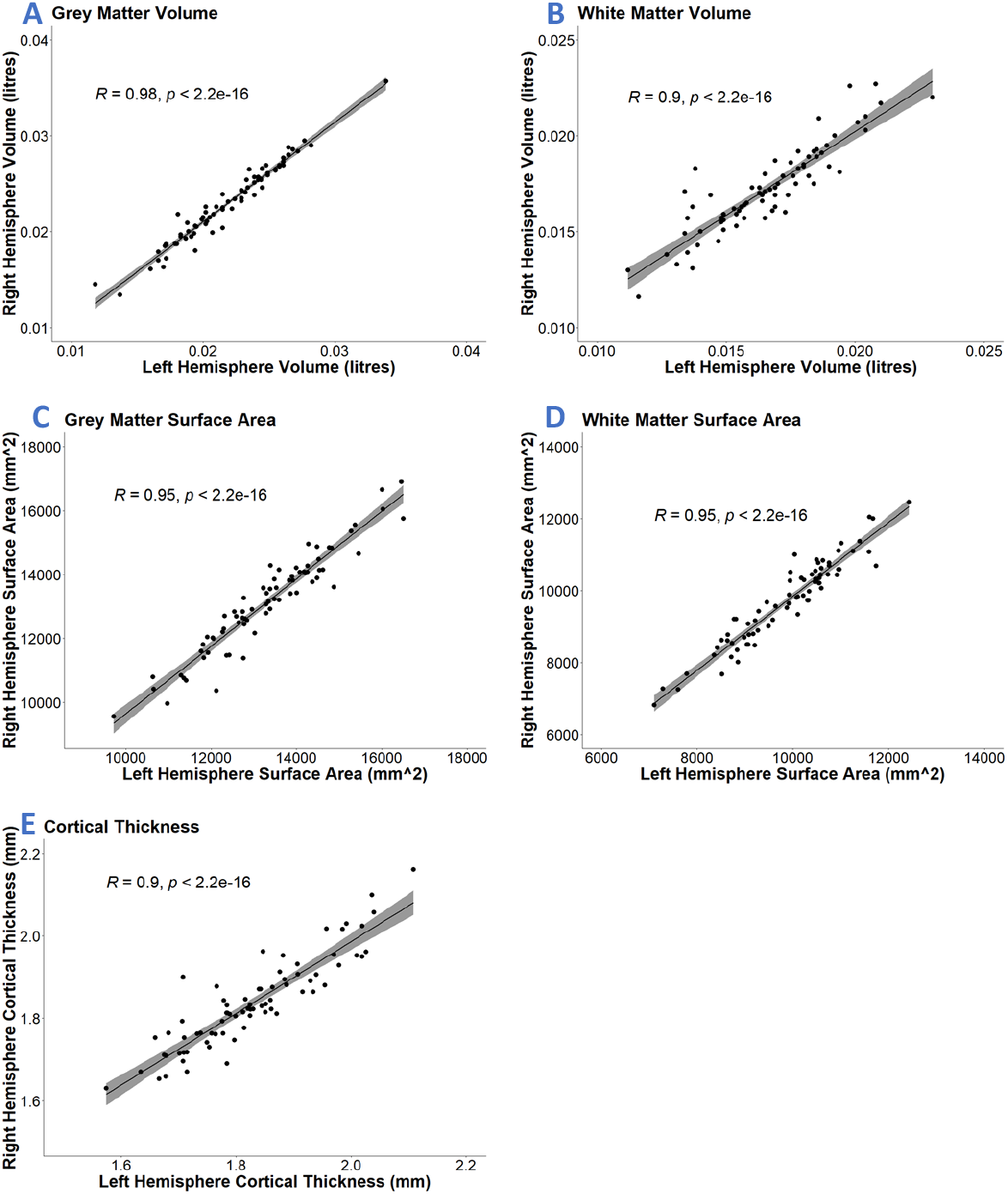
Hemisphere Comparison Graphs. Correlation between left and right hemisphere values for GM volume (A), WM volume (B), GM surface area (C), WM surface area (D) and cortical thickness (E). The linear fit and standard error are plotted, and R and p values are shown on each graph.

### 3.4. Scan-Rescan

To further evaluate the reliability of AutoMacq, the scan-rescan dataset was processed to give both volume-based and surface-based outputs. Over such a short span of time between ‘scan’ and ‘rescan’ (less than 1 week), noticeable structural brain changes are not expected. Rescan data was available for a subset of 13 Newcastle subjects (8 males and 5 females).

Results indicate strong correlations across metrics despite a modest sample size and the fact that the animals were scanned while awake (ICC values between 0.6 and 0.95, fig. 7).

**Figure 7:**
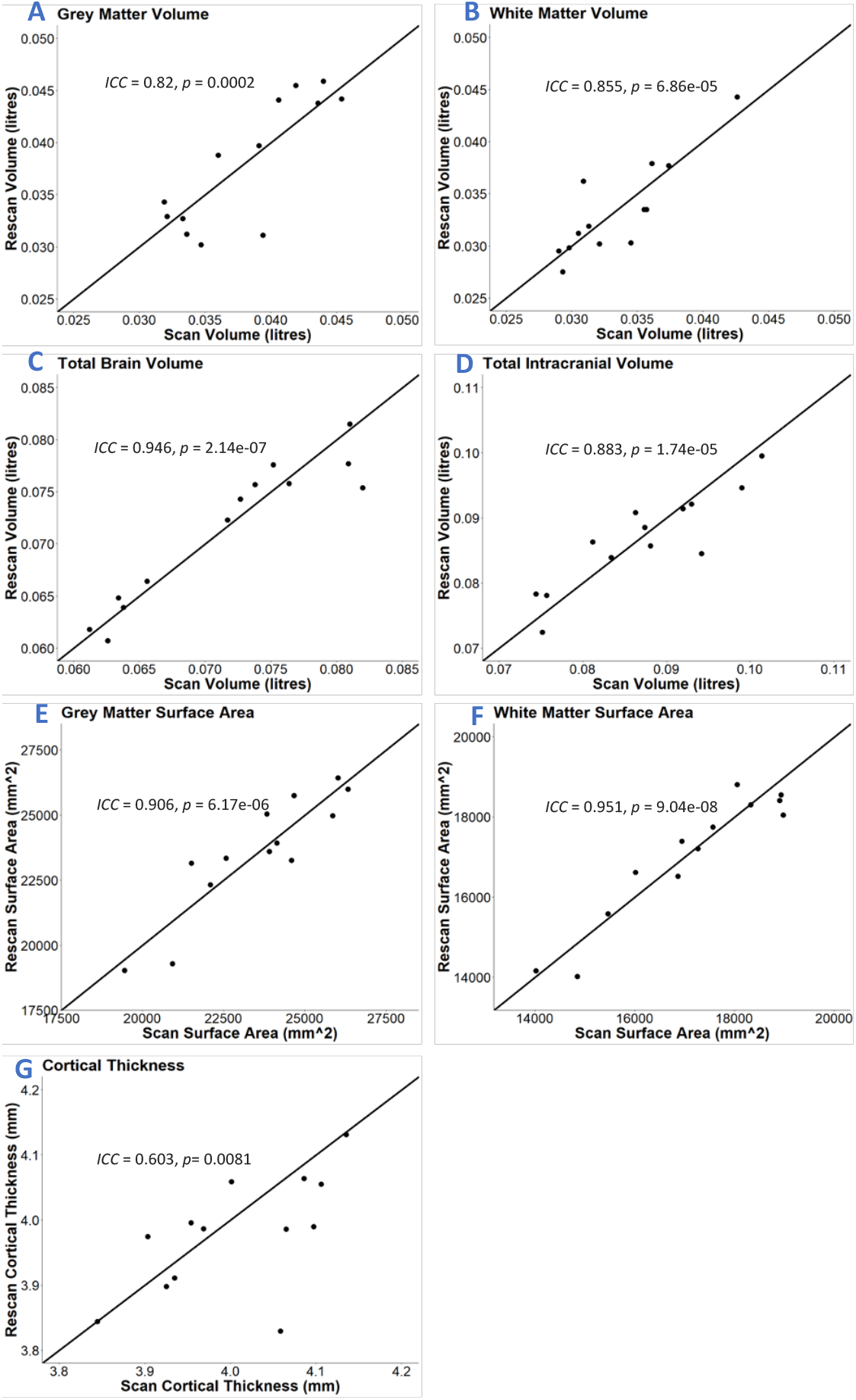
Scan-Rescan Graphs. Graphs of the correlation between the scans and the rescans for GM volume (A), WM volume (B), total brain volume (C), total intracranial volume (D), GM surface area (E), WM surface area (F) and cortical thickness (G). The identity line, ICC value and p value are shown on each graph.

### 3.5. Global Brain Changes with Ageing

To demonstrate a possible application of the AutoMacq pipeline, the impact of ageing on total GMV was tested using the male subjects from the cross-sectional datasets. Female subjects were excluded due to the small sample size, and the scans from any site with fewer than 5 male subjects were also excluded so that effect of site could be adequately controlled for in the model. A significant, linear decrease in total GMV with age was found (p=0.0066, fig. 8).

**Figure 8:**
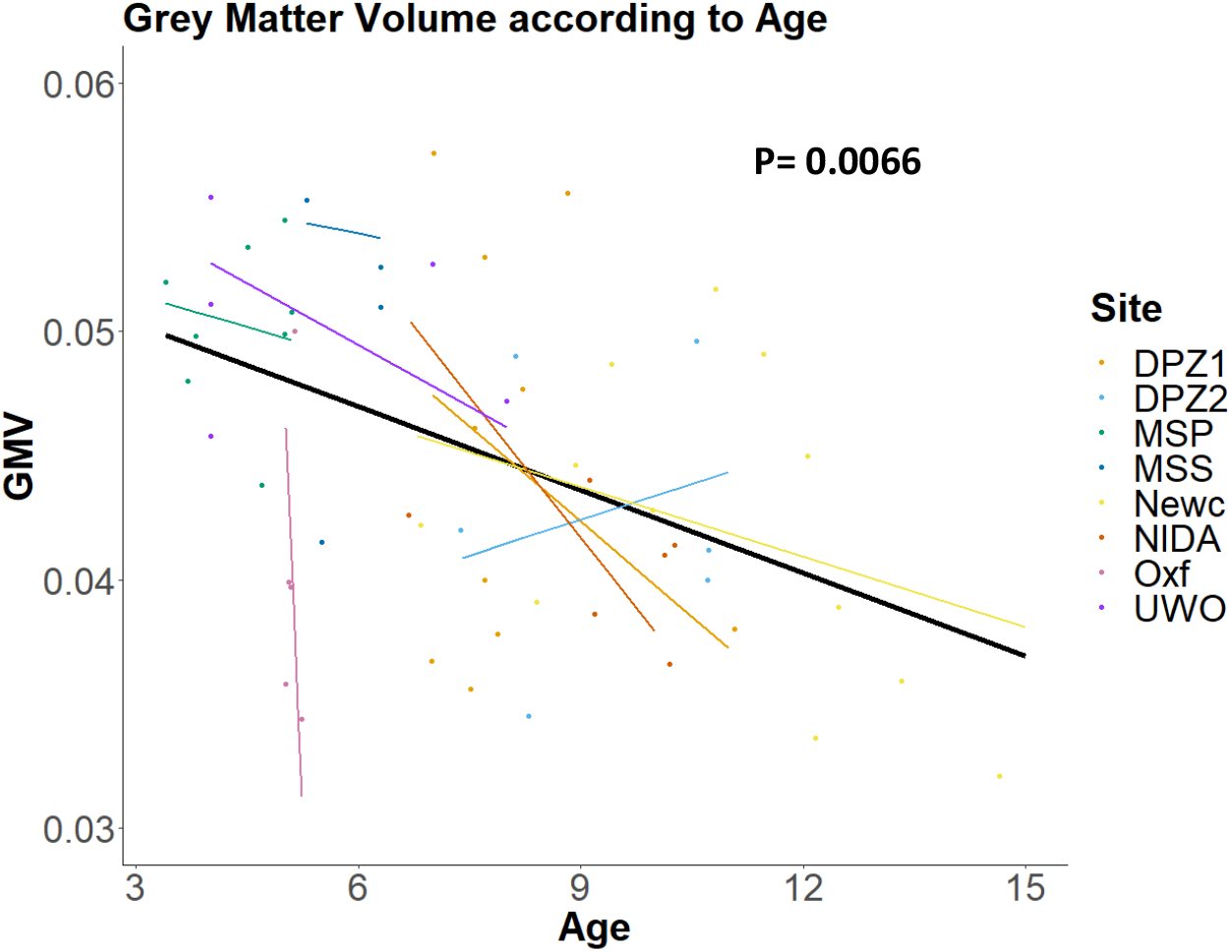
Changes in GM volume according to ageing. The bold black linear line corresponds to the main effect of age, while controlling for TIV and with site/scanner declared as a random effect. The thin coloured lines correspond to linear fits of age effect in each site while controlling for TIV. Dots correspond to raw data (unadjusted for TIV).

## 4. Discussion

### 4.1. Strengths of AutoMacq

AutoMacq is a robust processing pipeline, capable of successfully processing macaque MRI data with a wide range of quality and scan parameters, with minimal manual intervention. The two manual steps within the pipeline are simple to carry out and do not require any expert knowledge of macaque neuroanatomy. This, coupled with the automation of the rest of pipeline, makes AutoMacq relatively easy to use. Additionally, AutoMacq is unique amongst macaque pipelines in its ability to produce both voxel-based and surface-based metrics (Balbastre *et al*. 2017; Garcia-Saldivar et al. 2021; Lepage et al. 2021), allowing for more avenues of investigation and comparison with a wider range of previous studies (Goto *et al*. 2022).

Both T1 scans alone and datasets of T1 and T2 scans were successfully processed through AutoMacq. 100% of the scans processed through AutoMacq produced an accurate brain mask. This extremely high level of success in terms of brain extraction is better than the one obtained using the FSL bet function (Lepage et al. 2021) and comparable to what can be obtained by more sophisticated deep learning-based approaches (Wang et al., 2021). 97.3% of cross-sectional scans processed through AutoMacq gave good quality volume-based outputs and 87.8% gave good quality surface-based outputs. A good quality output was defined as one not requiring manual correction. The high percentage of good quality outputs produced illustrates AutoMacq’s accuracy, with fewer errors in the outputs from AutoMacq than those produced when using other pipelines to process data from various sites (Garcia-Saldivar *et al*. 2021; Lepage et al. 2021). This better performance is likely to come from the use of SPM segmentation routine to identify the GM/WM boundary. AutoMacq was able to handle scans with a wide range of scan parameters and quality, including those known to be difficult to process, such as scans acquired in awake subjects (Newcastle University dataset) and scans from subjects that have open skulls due to head implants (Milham et al., 2018; PRIMatE Data Exchange (PRIME-DE) Global Collaboration Workshop and Consortium 2020). The few scans for which AutoMacq produced outputs with errors were not all from one site, indicating that the poorer outputs were not due to an inability to handle specific scanning parameters but likely due to issues specific to the individual scans themselves.

The present assessment of AutoMacq’s quality as a processing pipeline is strengthened by the use of quantitative measures of reliability. The hemisphere comparison resulted in strong correlations for both volume-based and surface-based metrics, despite the fact that problematic volumes and surfaces were not excluded from the analyses. Since only minimal differences from biological origin were expected between hemispheres, this result indicates that AutoMacq produces reliable volume and surface outputs. The reliability of AutoMacq was then further demonstrated by the scan-rescan analysis. This analysis focused on a smaller sample (N=13) from the Newcastle dataset alone. Despite the limited sample size, good to excellent reliability was observed for all of the volume-based metrics tested and 2 of the 3 surface-based metrics, with the correlation for cortical thickness being weaker but still showing moderate reliability (Koo and Li 2016). This weaker correlation is unsurprising given how much influence sample size has in studies of cortical thickness (Pardoe *et al*. 2013), and the strength of this correlation did increase when the outlier was removed. Also, these correlations are fairly strong given the fact that macaques in this subset of data were all scanned whilst awake, and head movements are known to have a major impact on ICC in MRI studies (Hedges *et al*. 2022). Overall, this scan-rescan analysis provides further evidence for the strong reliability of the AutoMacq pipeline. Both the hemisphere comparison and scan-rescan analysis resulted in similar correlations to what has been seen for human MRI studies (Carmon *et al*. 2020; Hedges *et al*. 2022).

A key strength of AutoMacq is the ability to carry out both VBM and SBM. This is an advantage as it allows for data to be exploited in multiple different ways, and the optional ability to substitute a macaque parcellation schema into the FreeSurfer processing compounds this. Furthermore, the ability to produce both volume-based and surface-based metrics allows for comparison to a wider range of studies, which is particularly important due to the continued publication of both VBM and SBM human studies, especially in clinical populations (Goto *et al*. 2022). Historically, human cortical VBM analyses have been criticised because they tended to suffer from volumetric projection to a template, particularly due to the highly variable cortical folding pattern between subjects, and SBM has been - in part-developed to avoid these issues (Postelnicu *et al*. 2008; Villalon *et al*. 2011). However, the cortical folding pattern in macaques is much more preserved between subjects (Van Essen *et al*. 2019). Additionally, the probability of problems linked to partial volume effects can be mitigated by the use of high magnetic field strengths, allowing the acquisition of images at higher spatial resolution (Milham *et al*. 2018). We therefore suspect that the potential drawbacks of VBM in human data are less likely to be relevant for macaque analyses, which could be tested in future using the AutoMacq pipeline.

### 4.2. Limitations of AutoMacq

One limitation of AutoMacq is our use of a human atlas for the FreeSurfer processing. This is what necessitates the manual correction of the atlas registration. The inability to run AutoMacq fully automatically may make processing very large datasets more time consuming, however datasets of hundreds or thousands of macaque MRI scans are relatively rare currently. Additionally, AutoMacq produced good outputs for the vast majority of subjects despite the use of a human atlas. It is possible that substituting in a macaque atlas may result in even fewer errors but given the high success rate already observed this is likely to be unnecessary for most studies.

### 4.3. Impact of Ageing on GMV

The impact of ageing on total GMV was investigated using 59 male subjects aged between 3 and 15 years, scanned using 8 different scanners from across 7 different sites. Given macaques age at 4 times the rate of humans during childhood (reaching sexual maturity around 4 and full physical maturity around age 5-6) and 3 times the rate of humans during adulthood (Mattison and Vaughan 2017), the age range tested here can be roughly compared to ages of 12-45 in humans.

A significant linear decrease in GMV was found, indicating that in male rhesus macaques there is a significant decline in GMV prior to reaching mid adulthood. This is a novel finding for this age group and suggests that the age-related decrease in GMV seen in mid/late adulthood in previous studies (Wisco *et al*. 2008; Chen *et al*. 2013) may actually start earlier in the lifespan. This finding fits with what has been seen in human studies, where a decline in GMV has been seen to occur across adolescence and early adulthood (Bartzokis *et al*. 2001; Lebel *et al*. 2012, Bethlehem *et al*. 2022). This study therefore provides further strength to rhesus macaques being models of healthy human ageing (Phillips *et al*. 2014; Roefsema and Treue 2014; Stonebarger *et al*. 2021).

It should be noted that this study only utilised male subjects, as scans were only available from very few females, and it would not have been possible to control for sex. It is therefore not possible to fully generalise these results to female rhesus macaques.

## 5. Conclusion

AutoMacq offers a processing pipeline for rhesus macaque MRI data that is easy to use and can be completed without expert knowledge of macaque neuroanatomy. AutoMacq can process data with a wide range of quality and parameters, from across different sites and scanners, with a high level of success. The pipeline is unique amongst macaque processing pipelines in its ability to generate both surface-based and voxel-based metrics, offering two ways to exploit macaque MRI scans and allowing for easier comparison to a wider range of previous research.

## 6. Acknowledgements

With thanks to Dr Dirk Jan Ardesch for his guidance on FreeSurfer and his assistance in writing the initial FreeSurfer scripts. Thank you to Professor Hank P. Jedema and Dr Charles W Bradberry for sharing the NIDA dataset, Dr Jerome Sallet for sharing the Oxford dataset, and the Cognitive Neuroscience Laboratory at the German Primate Center for sharing the DPZ dataset.

