## Supplementary Materials for "AutoMacq: an automatic pipeline to analyse macaque structural MRI data"

**Supplementary Data- Materials and Methods**

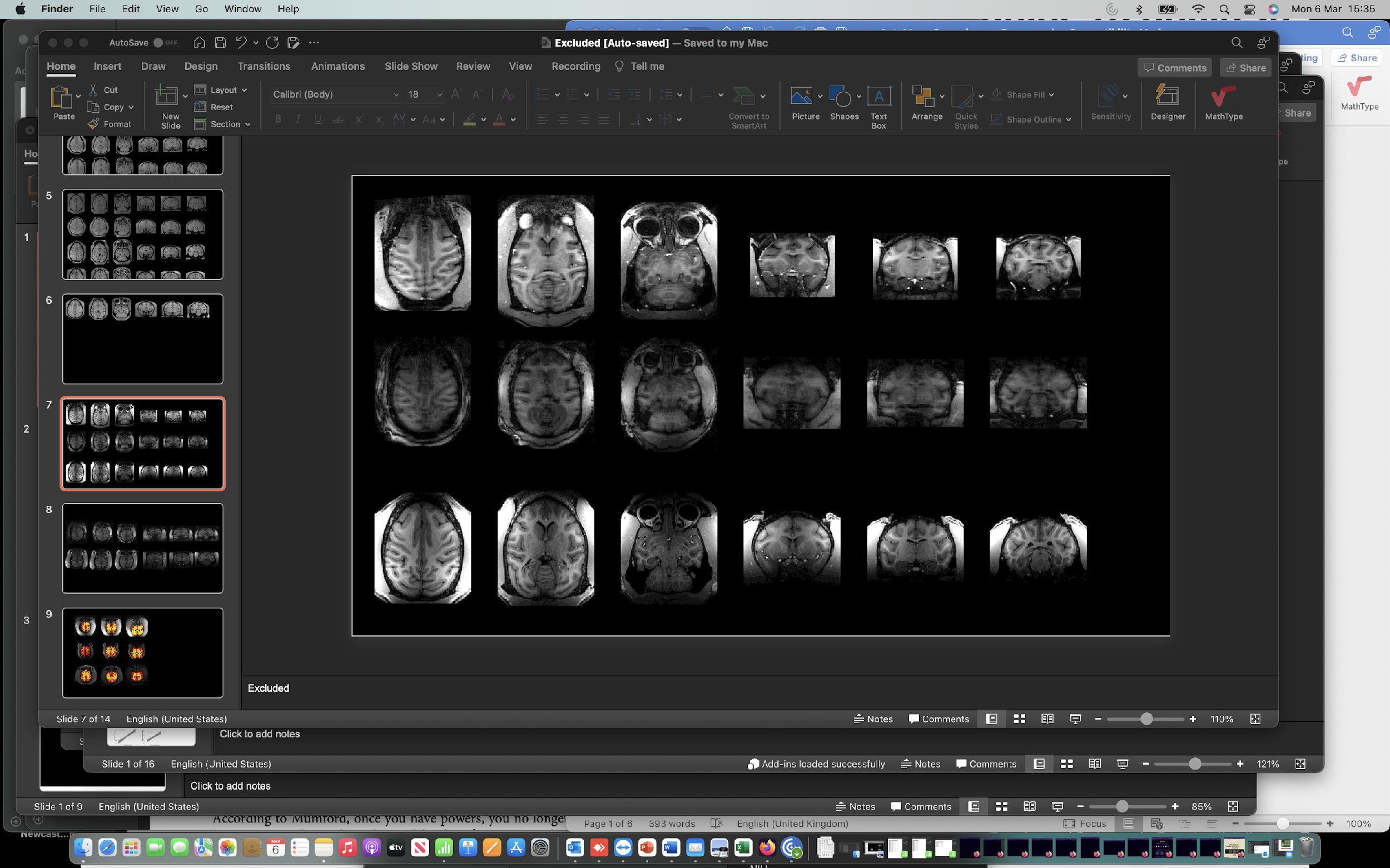

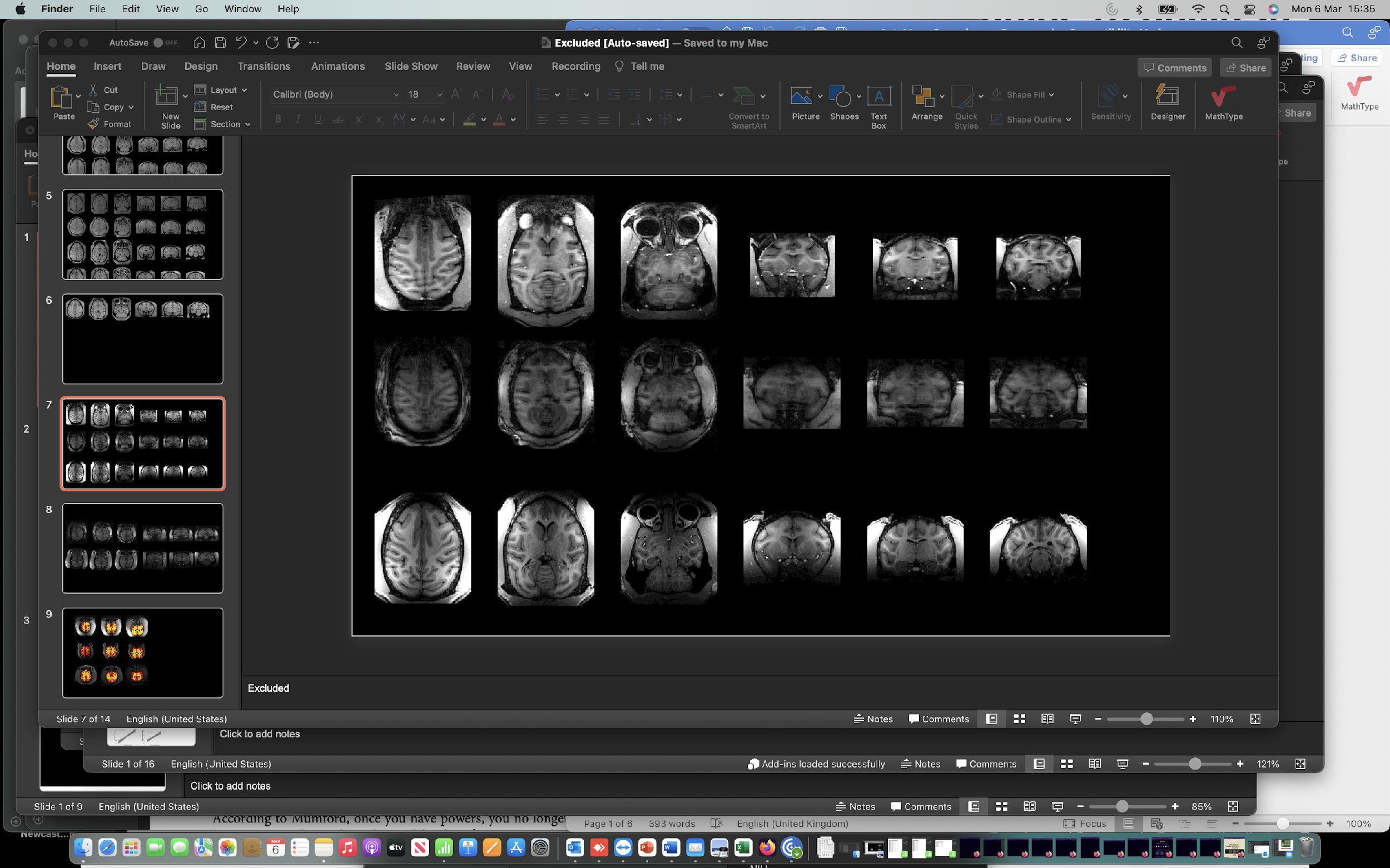

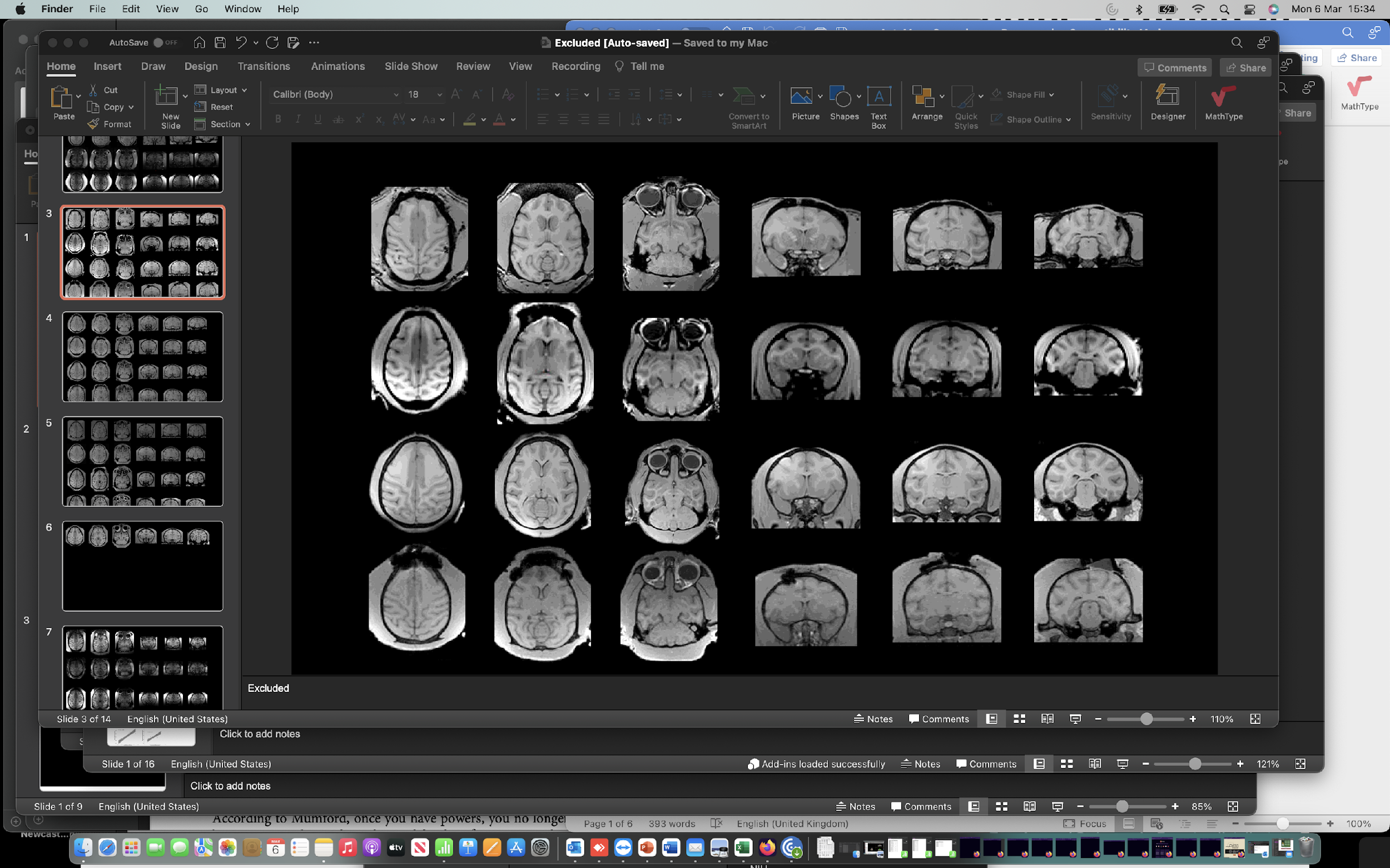

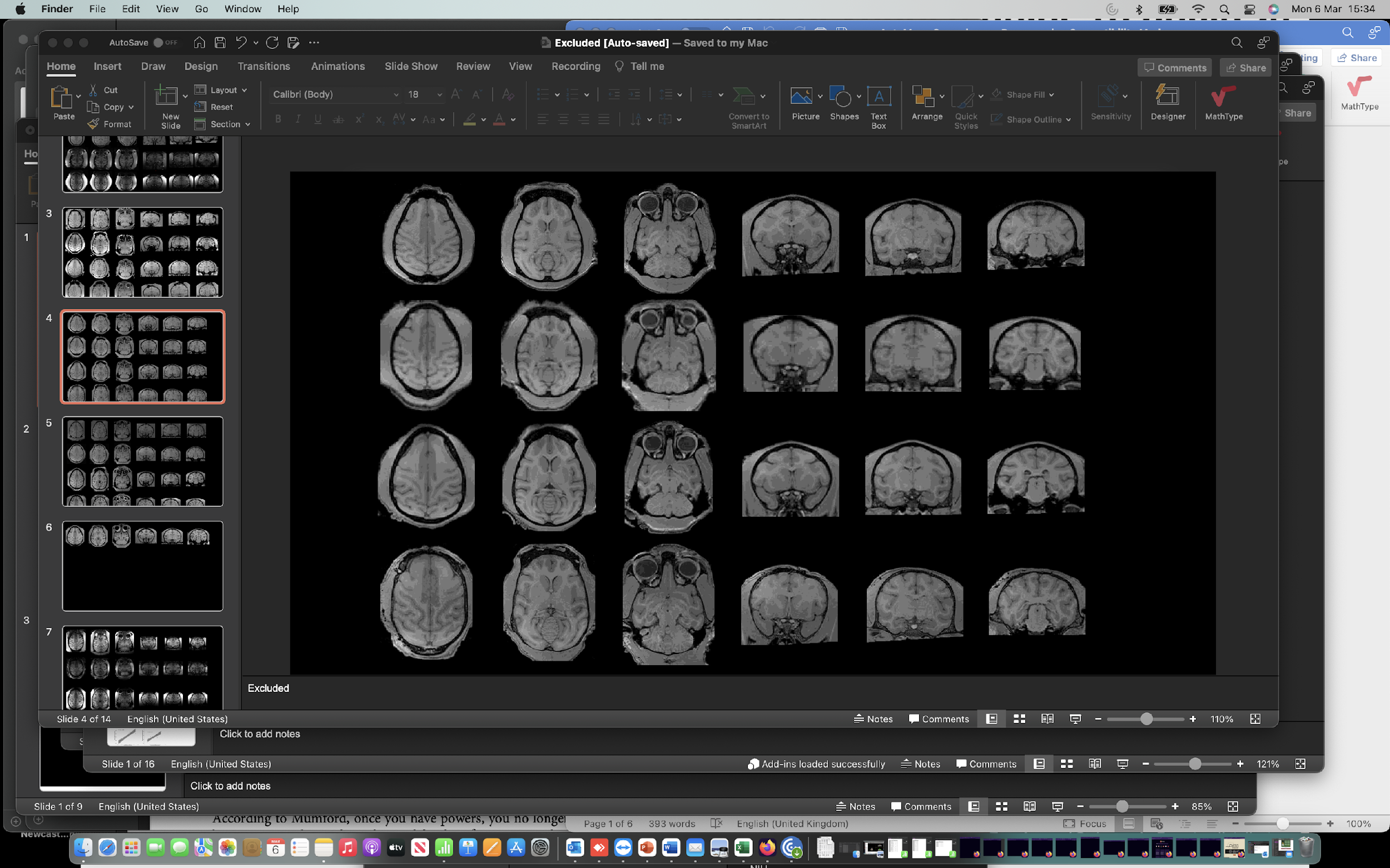

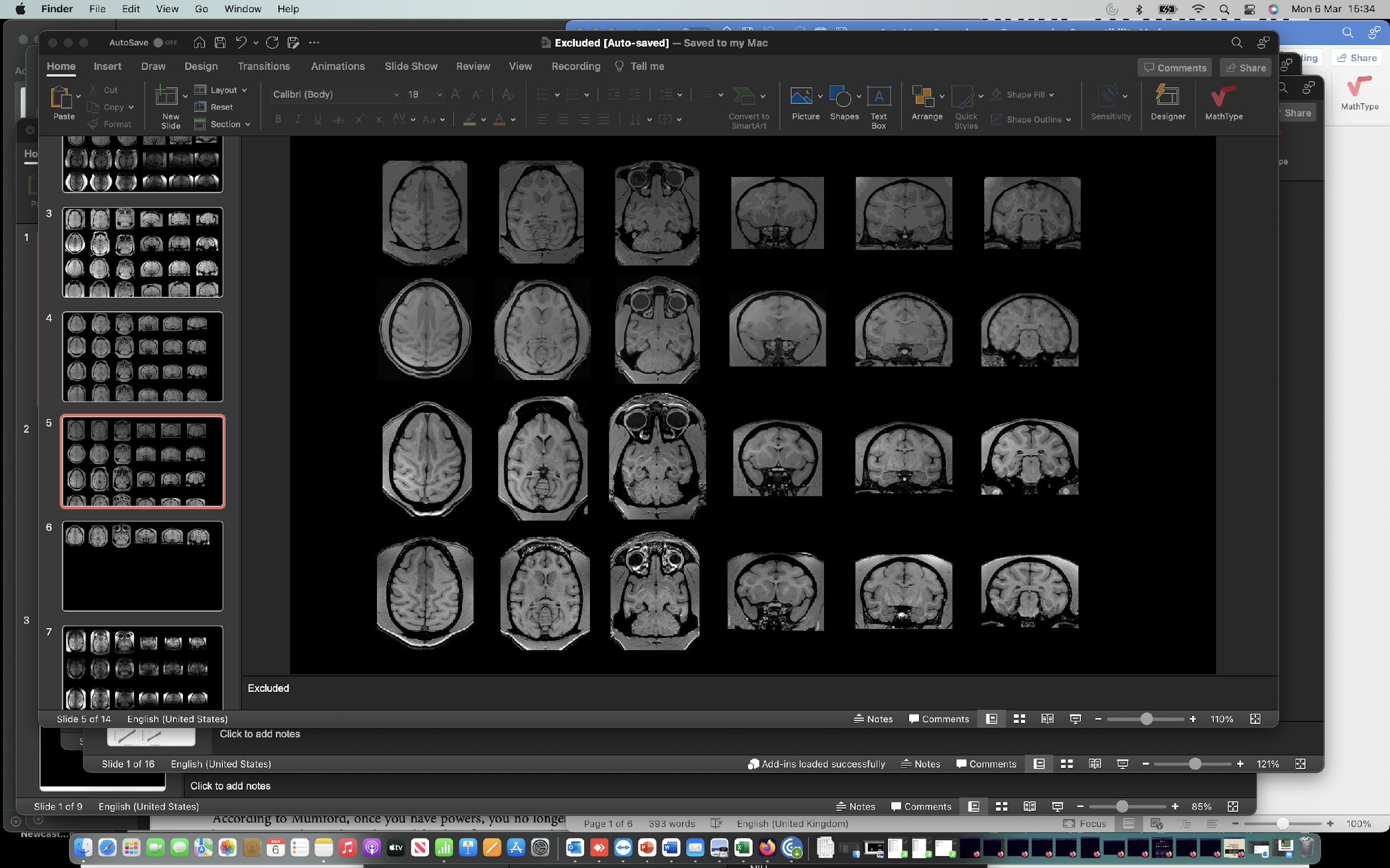

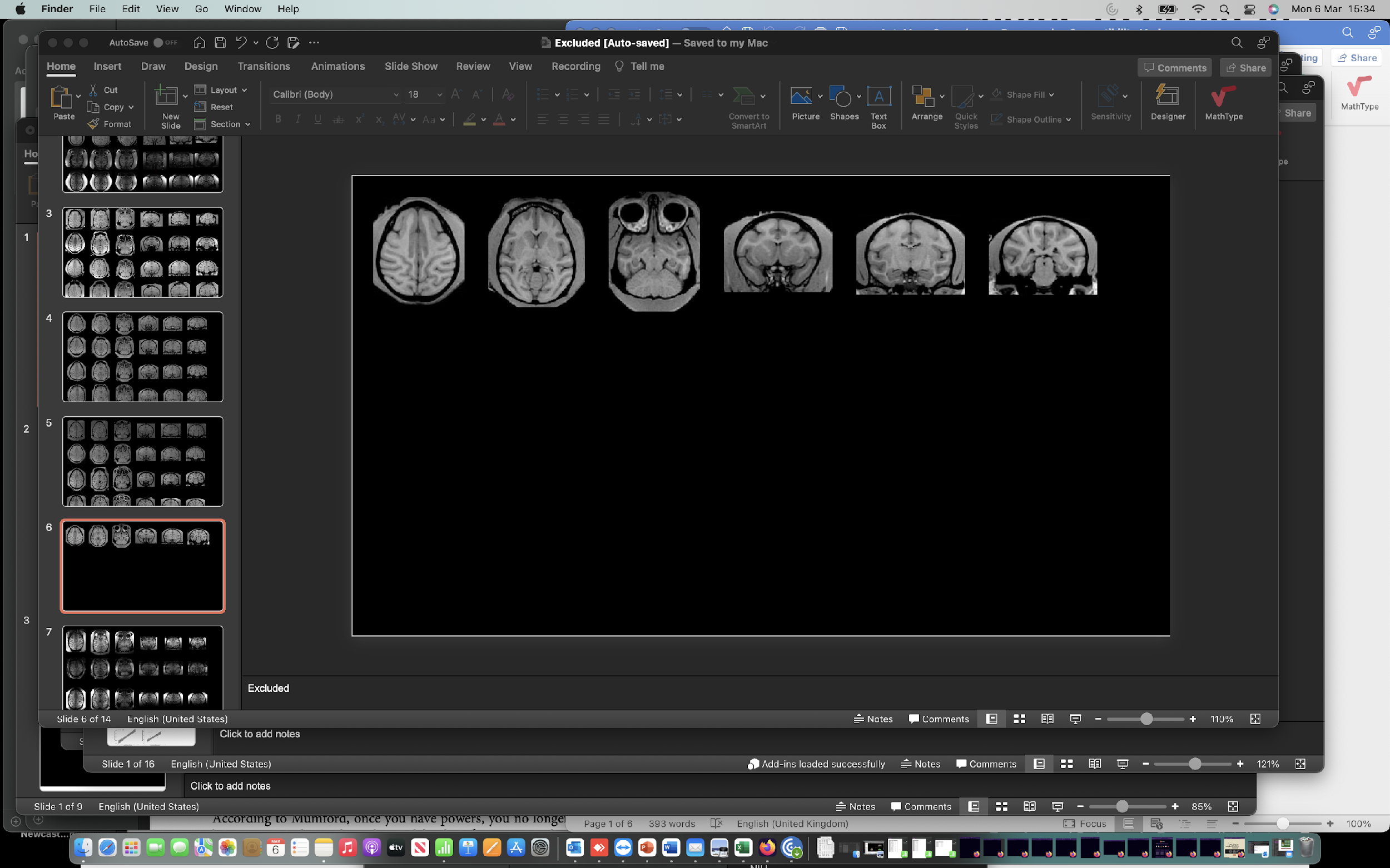

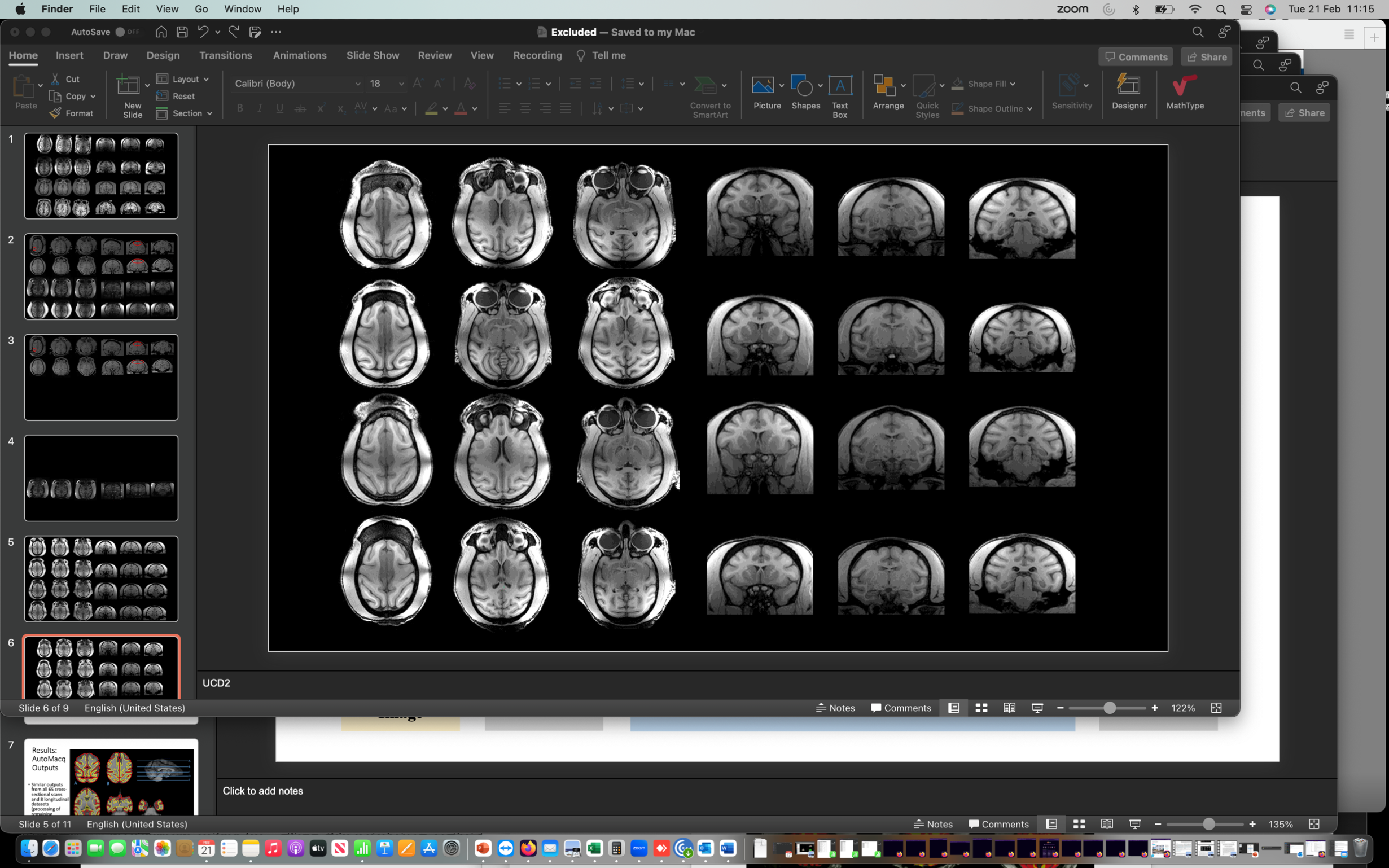

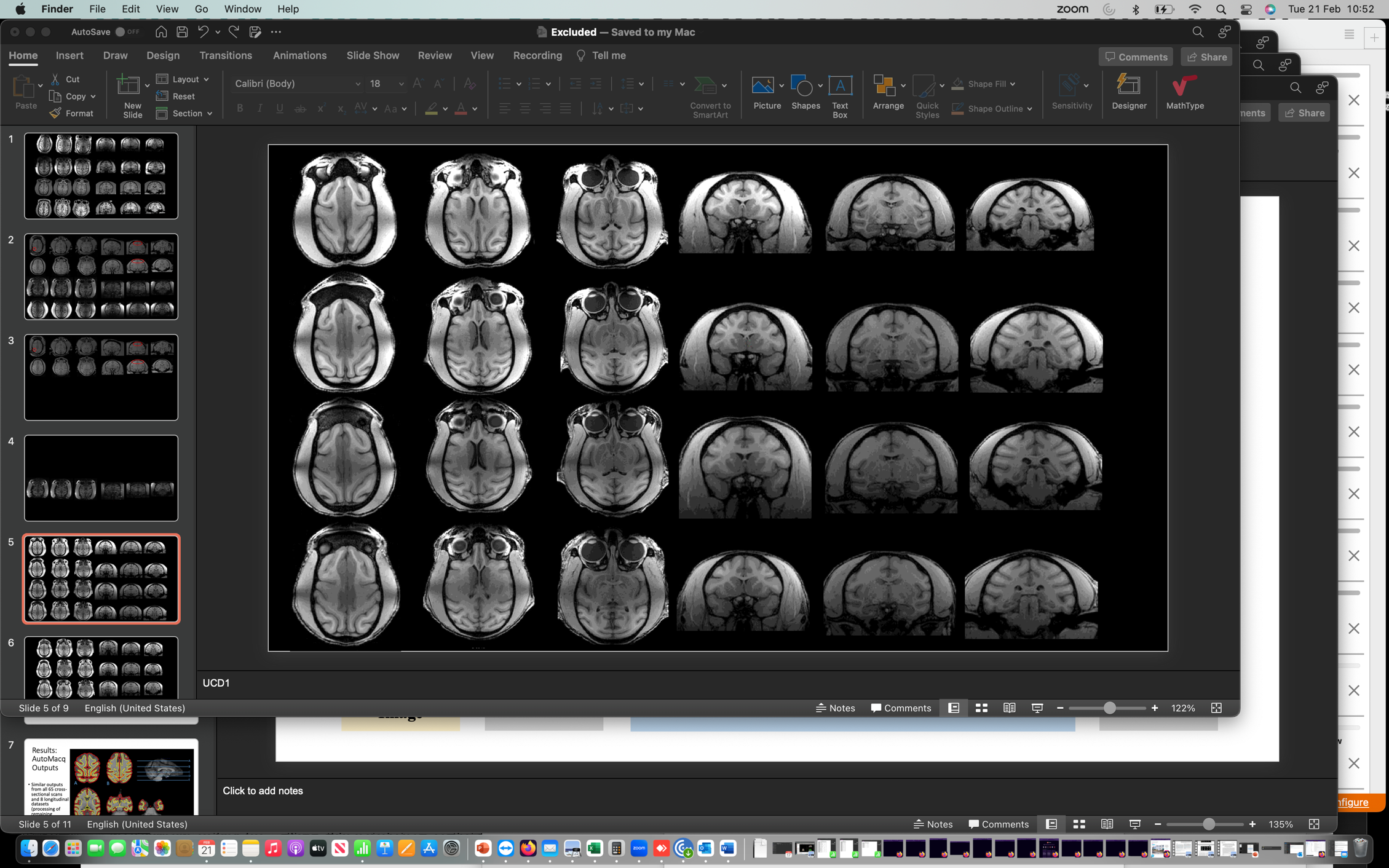

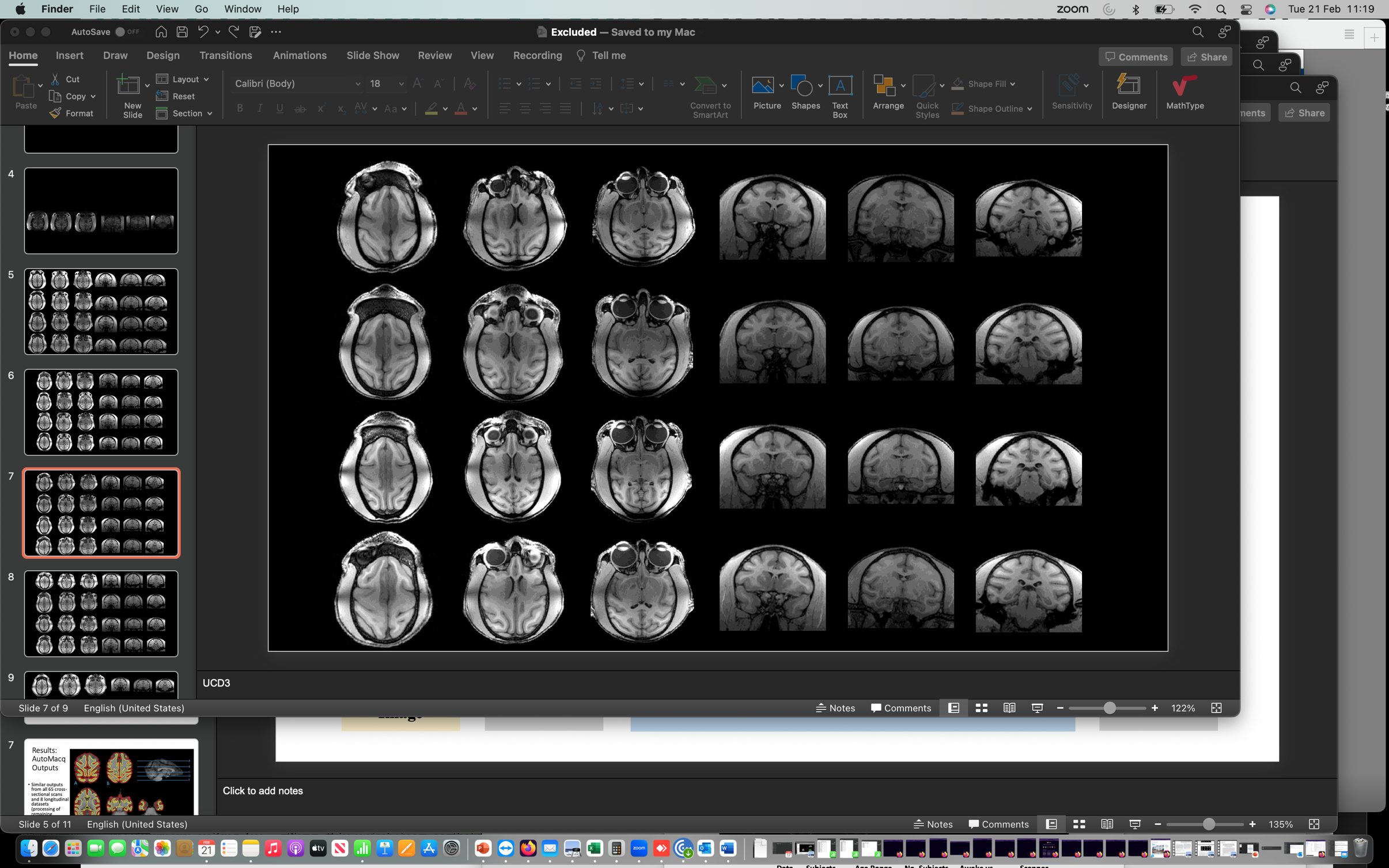

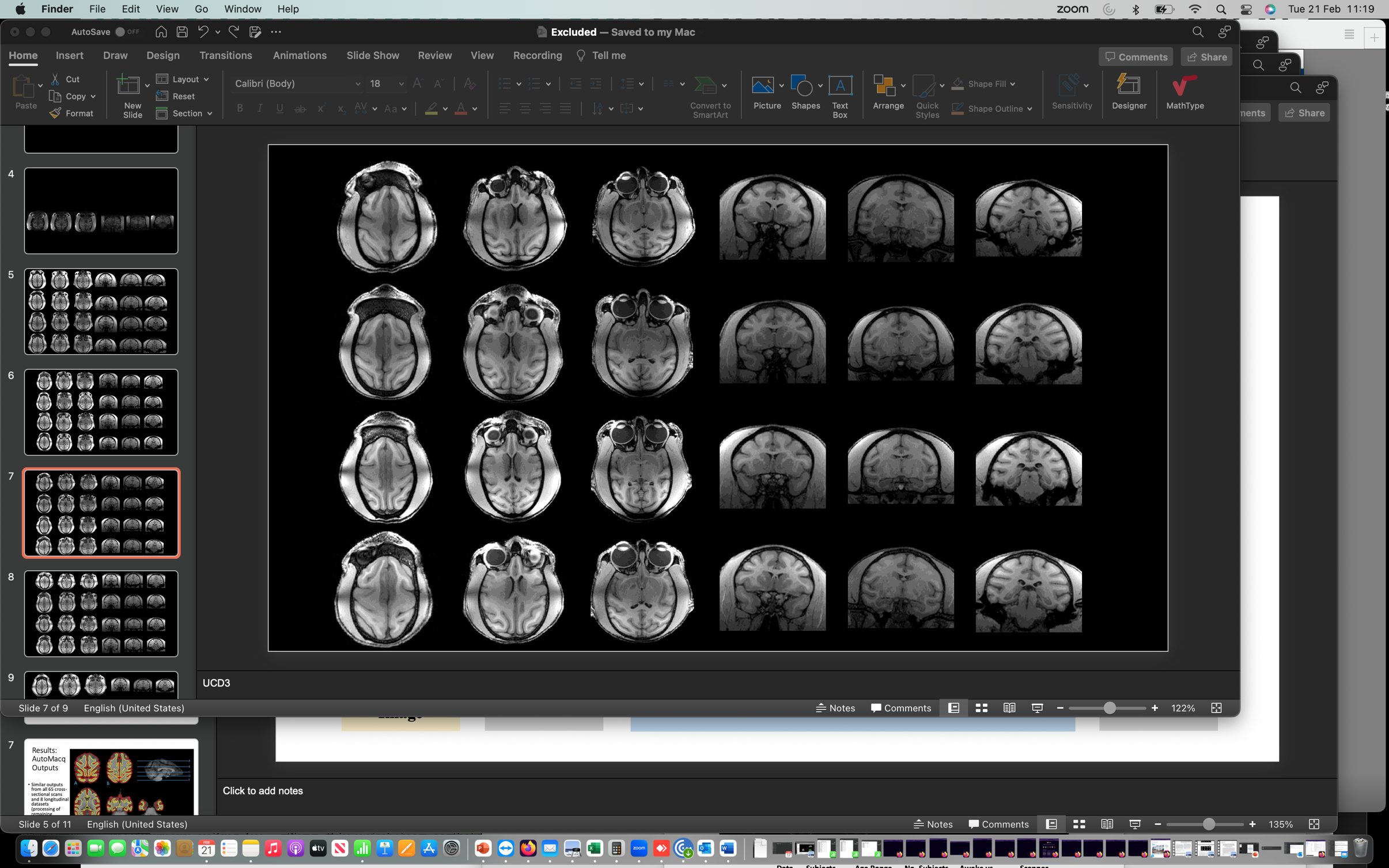

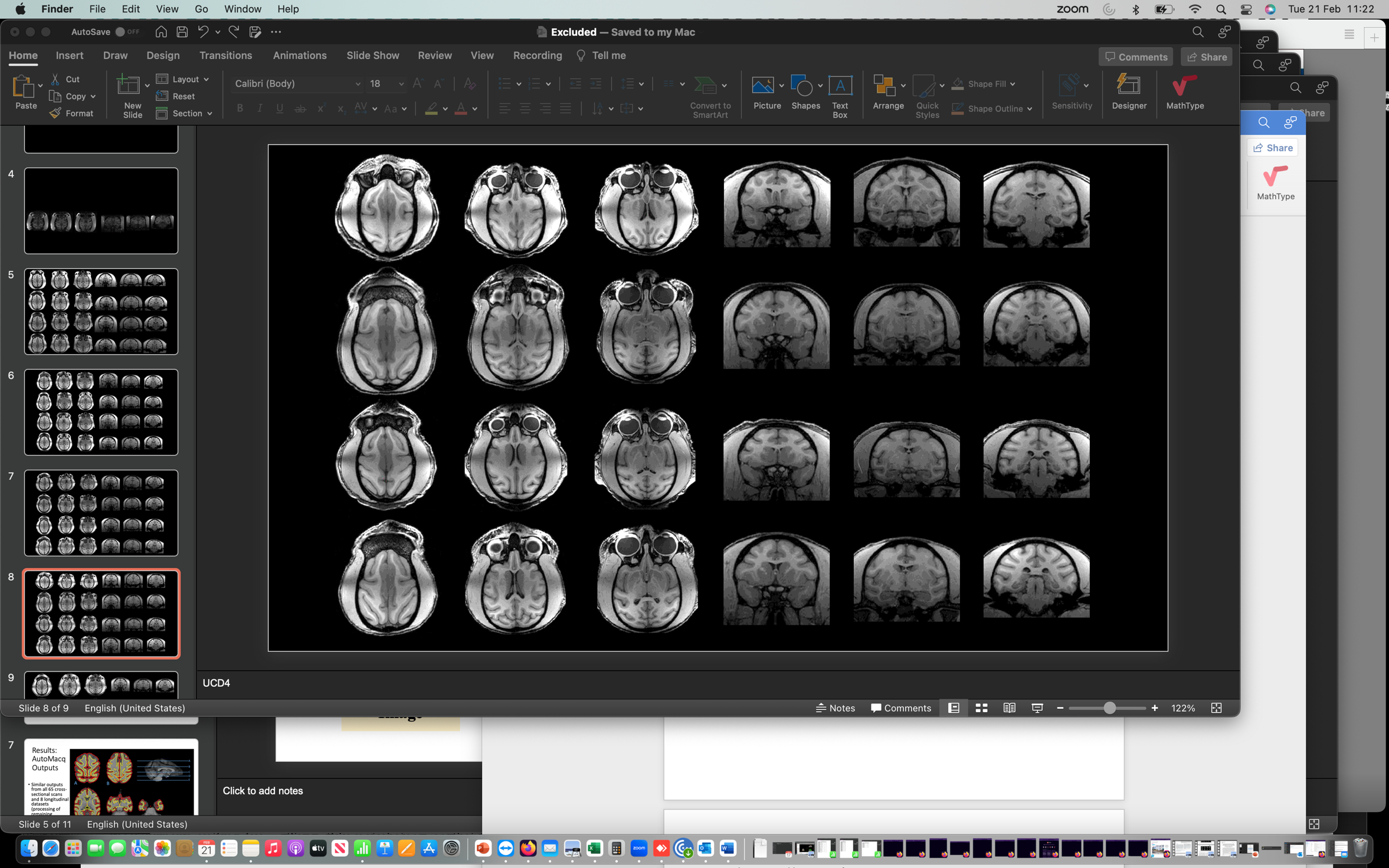

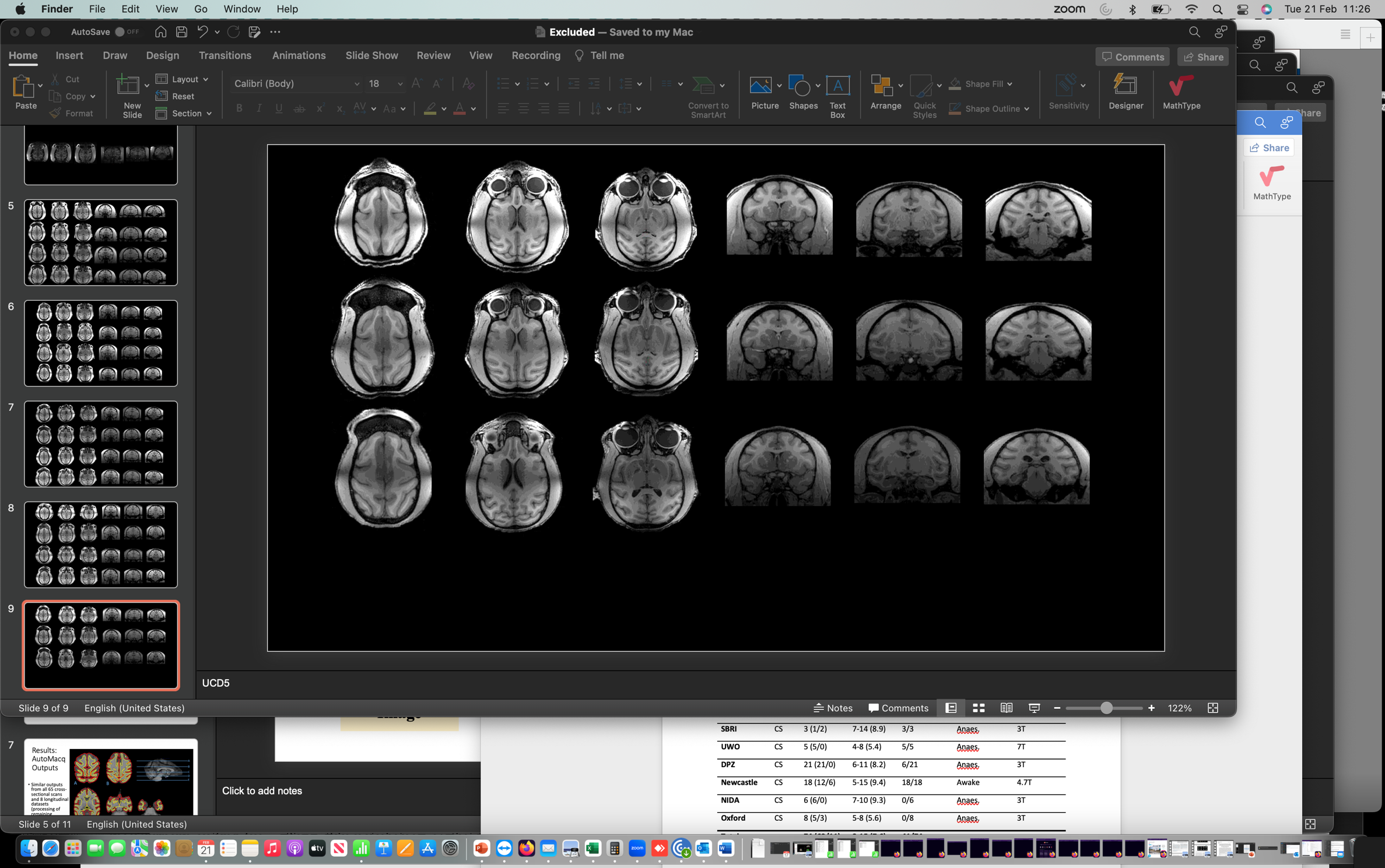

**Supplementary Figure 1- Slices from Excluded Scans.** 3 horizontal and 3 coronal slices from the scans of every subject excluded due to image quality issues. The subjects are separated by site (A= Newcastle, B= Oxford, C= DPZ, D= UC-Davis) and areas where issues are particularly prominent are highlighted. One subject was excluded due to motion artefacts (A2), 2 subjects were excluded due to prominent hyperintensities (A1 and B1), and the remaining ones (n=32) were excluded due to poor contrast to noise ratio in some parts of the brain.

**Supplementary Table 1- Detailed scan parameters for each site**

|  | **Scanner Type** | **Coils Used** | **Sequences Used** | **Voxel Size** | **TE**  **T1 (T2)** | **TR**  **T1 (T2)** |
| --- | --- | --- | --- | --- | --- | --- |
| **Newcastle** | Bruker 4.7T vertical | 4 channel parallel imaging coils | MPRAGE, RARE | 0.608 x 0.608 x 0.618mm | 3.75ms (14.33ms) | 2000ms (12296.7ms) |
| **DPZ1** | Siemens Prisma 3T |  |  | 0.5 x 0.5 x 0.5mm | 2.96ms | 2700ms |
| **DPZ2** | Siemens Prisma 3T? |  |  | 0.5 x 0.5 x 0.5mm |  |  |
| **DPZ3** | Siemens Magnetom Trio |  |  | 0.65 x 0.651 x 0.651mm |  |  |
| **DPZ4** | Siemens Magnetom TIM Trio |  |  | 0.5 x 0.5 x 0.5mm |  |  |
| **NIDA** | Siemens 3T Allegra | Custom-designed holder/secondary coil (NOVA_16) | MPRAGE | 0.6 x 0.598 x 0.598mm | 3.04ms | 1680ms |
| **Oxford** | 3T whole body scanner | 4-channel, phased-array, radio-frequency coil w/ local transmission coil | MPRAGE | 0.5 x 0.5 x 0.5mm | 4.01ms | 2500ms |
| **MSP** | Philips Achieva 3T | 4-channel phased array coil (Windmiller-Kolster Scientific), transmit through body coil |  | 0.5 x 0.5 x 0.5 mm | 6.93ms (366ms) | 1500ms (2500ms) |
| **MSS** | Siemens Skyra 3T | 4-channel clamshell coil | MPRAGE | 0.5 x 0.5 x 0.5 mm | 3.02ms (539ms) | 2700ms (3200ms) |
| **SBRI** | Siemens Prisma 3T | 2 x L11 and 1 x L7 Siemens ring coils | T1 MPR 3D, T2 SPACE TRA | 0.5 x 0.5 x 0.5 mm | 3.62s (366ms) | 3000ms (3000ms) |
| **UWO** | Siemens Magnetom 7T | Custom-made 24-channel phased array receive coil w/ 8-channel transmit coil | MPRAGE | 0.5 x 0.5 x 0.5 mm | 3.88ms | 65000ms |

TE: Echo Time, TR: Repetition Time

**Supplementary Text- Longitudinal Processing**

Longitudinal studies benefit from a focus on within-subject differences, removing any impact of genes or gene x environment interactions and increasing statistical power (Song *et al.* 2021). For MRI studies, pipelines which handle cross-sectional data are not fully leveraging the potential of longitudinal data, as they are optimised to generalise across a cohort, often at the cost of within-individual accuracy. For example, in a longitudinal pipeline, subject-specific templates could be used to improve various steps of the pipeline.

AutoMacq is intrinsically capable of processing longitudinal MRI datasets from rhesus macaques. The adaptations to the cross-sectional pipeline required to do so are outlined below.

**Adaptations to AutoMacq- Longitudinal Voxel-based Processing**

The stages to obtain voxel-based metrics for longitudinal datasets using AutoMacq are very similar to those used for cross-sectional data, with one additional step. After all the scans for a subject have been cropped and reoriented (as described in section 2.3.1), a longitudinal registration step is performed. This includes a rigid-body and a diffeomorphic registration of each T1 scan to a mid-point average image (Ashburner and Ridgway, 2013). If T2 data are available, a longitudinal registration of T2 scans is carried out as well, and the average T2 scan is then coregistered to the average T1 scan. The rest of the voxel-based processing in AutoMacq is carried out in the same way it would be for individual cross-sectional scans but using the average image(s) as input file(s).

**Adaptations to AutoMacq- Longitudinal Surface-based Processing**

The standard FreeSurfer longitudinal processing stream for human data involves 3 major stages: 1) cross-sectional processing of all timepoints for each subject, 2) creation and processing of an unbiased template for each subject, 3) Processing of all timepoints for each subject using the subject-specific template. The longitudinal processing with AutoMacq therefore starts with the cross-sectional processing of all timepoints, as described in section 2.3. This cross-sectional processing provides a normalised, skull-stripped, atlas-registered T1 scan for every timepoint, which are then used to create a subject-specific template using robust, inverse consistent registration (Reuter et al., 2010). This template is used to help at various stages of the longitudinal re-processing of all time points: for example, skull stripping, Talairach transforms, atlas registration as well as spherical surface maps and parcellations are initialized with common information from the within-subject template, significantly increasing reliability and statistical power (Reuter et al., 2012).

The third stage is the actual longitudinal processing of the time points and is unchanged from the standard FreeSurfer longitudinal processing carried out for human MRI data. Briefly, every timepoint is processed again, but the template created in the previous step is used to initialise processing steps involved in the cortical and subcortical segmentation. This should increase the robustness and sensitivity of the overall longitudinal analysis.

**Testing AutoMacq Longitudinal Processing**

Longitudinal datasets from 3 different sites were available: Newcastle, NIDA and DPZ (see Suppl. Table 1). 3 to 13 scans were available for each subject (96 scans in total), with consecutive scans separated by at least 3 months.

**Supplementary Table 2: Description of Longitudinal datasets**

| **Site** | **Included Subjects**  **(M/F)** | **Age Range (in years)** | **Subjects with T2 data** | **Awake vs. Anaes.** | **Scanner Strength** |
| --- | --- | --- | --- | --- | --- |
| DPZ | 1 (1/0) | 6-8 | 0 | Anaes. | 3T |
| Newcastle | 10 (8/2) | 6-15 | 10 | Awake | 4.7T |
| NIDA | 5 (5/0) | 6-10 | 0 | Anaes. | 3T |
| *Total* | *16 (14/2)* | *6-15* | *10* |  |  |

M: male and F: female. Anaes.: anaesthetised.

For longitudinal VBM processing in AutoMacq, 87.5% of subjects (14/16) produced tissue volumes of similar quality to figure 2, and 12.5% of subjects showed errors in the segmentation of grey and white matter (2/16).

For longitudinal SBM processing in AutoMacq, the total 96 scans from the 16 subjects were processed through FreeSurfer. 59.4% (57/96) of scans gave good quality surfaces, and 40.6% of scans (39/96) showed errors in their surfaces. For both the VBM and SBM processing, errors were limited to subjects from Newcastle University. Supplementary figure 1 provides a comparison of surfaces produced using cross-sectional versus longitudinal processing, for the same scan, where there was some improvement with longitudinal processing.

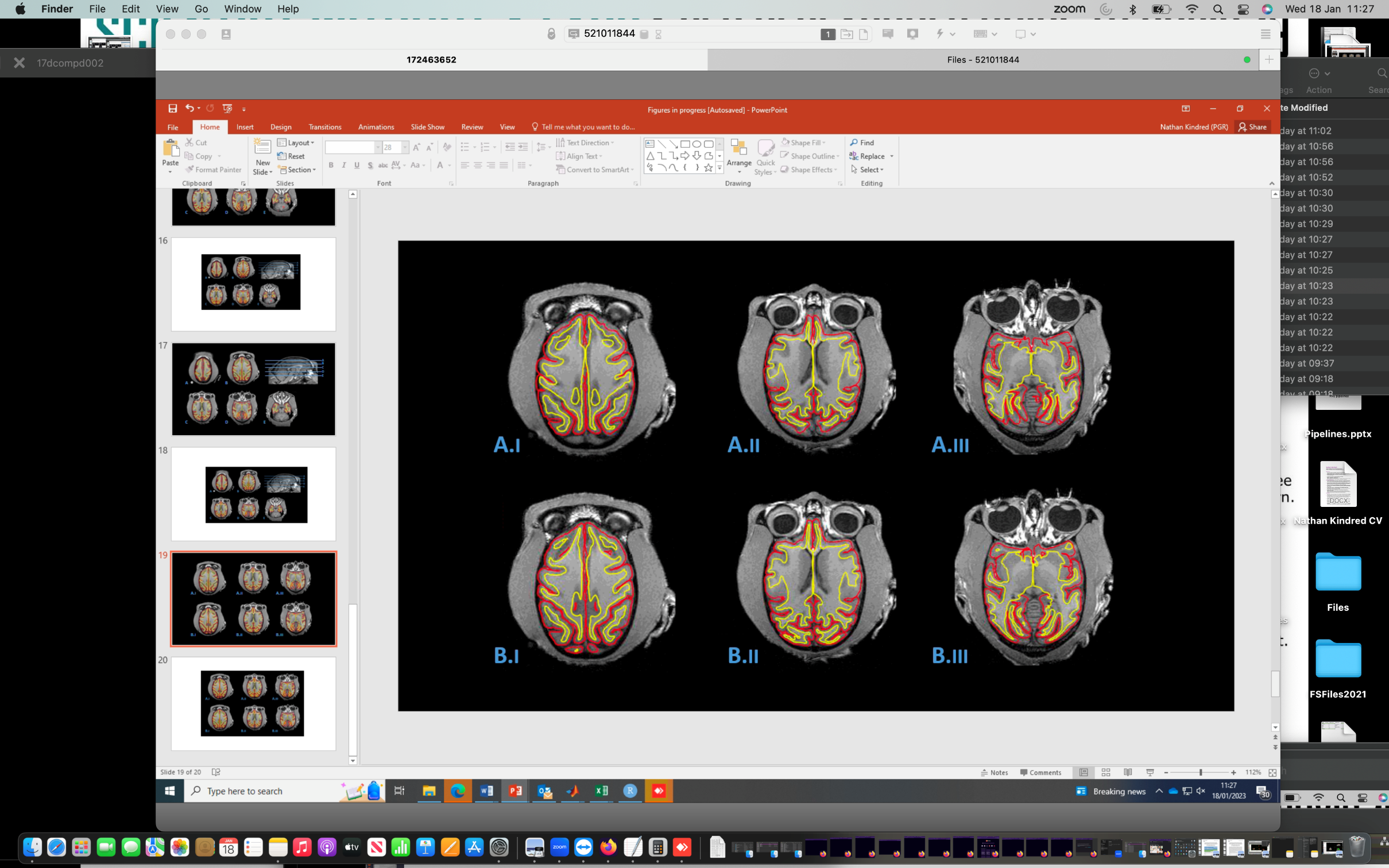

**Supplementary figure 2:** **Comparison of surfaces produced using the cross-sectional vs longitudinal AutoMacq pipeline.** Horizontal slices of examples of pial (shown in red) and white matter (shown in yellow) surfaces, produced for the same subject using the cross-sectional AutoMacq pipeline (A) and using the longitudinal AutoMacq pipeline (B).

**Discussion- Longitudinal Processing**

For voxel-based morphometry, AutoMacq had a comparable success rate for cross-sectional and longitudinal datasets, with good quality outputs (defined as those not requiring manual correction) being produced for 95.9% of cross-sectional scans and 87.5% of longitudinal subjects. This was despite the far larger sample size for the cross-sectional study (74 scans compared to just 16 longitudinal subjects). In terms of surface-based morphometry, the error rate for AutoMacq was higher with longitudinal data (40.6%) compared to with cross-sectional data (12.2%). It should be noted that almost a third of the subjects with longitudinal data (5/16) were part of the group of subjects which produced surface errors in the cross-sectional study. Because of the way the FreeSurfer longitudinal stream is carried out, errors in the cross-sectional processing of any single timepoint will be transferred to the template, inducing inaccuracies in the re-processing of each timepoint using the subject-specific template, providing a potential explanation for the increase in errors seen with AutoMacq longitudinal processing. Also, the longitudinal data available for testing AutoMacq was comprised mainly of subjects from Newcastle University which were both scanned awake and had open skulls due to head implants. This likely impacted on the image quality which may have also contributed to the increased proportion of errors for the longitudinal processing, as errors only occurred for subjects from the Newcastle dataset. For the subjects from other sites, which were scanned whilst anaesthetised, all of their scans were successfully processed through AutoMacq without errors for either VBM or SBM outputs.

**Supplementary Data- Results**

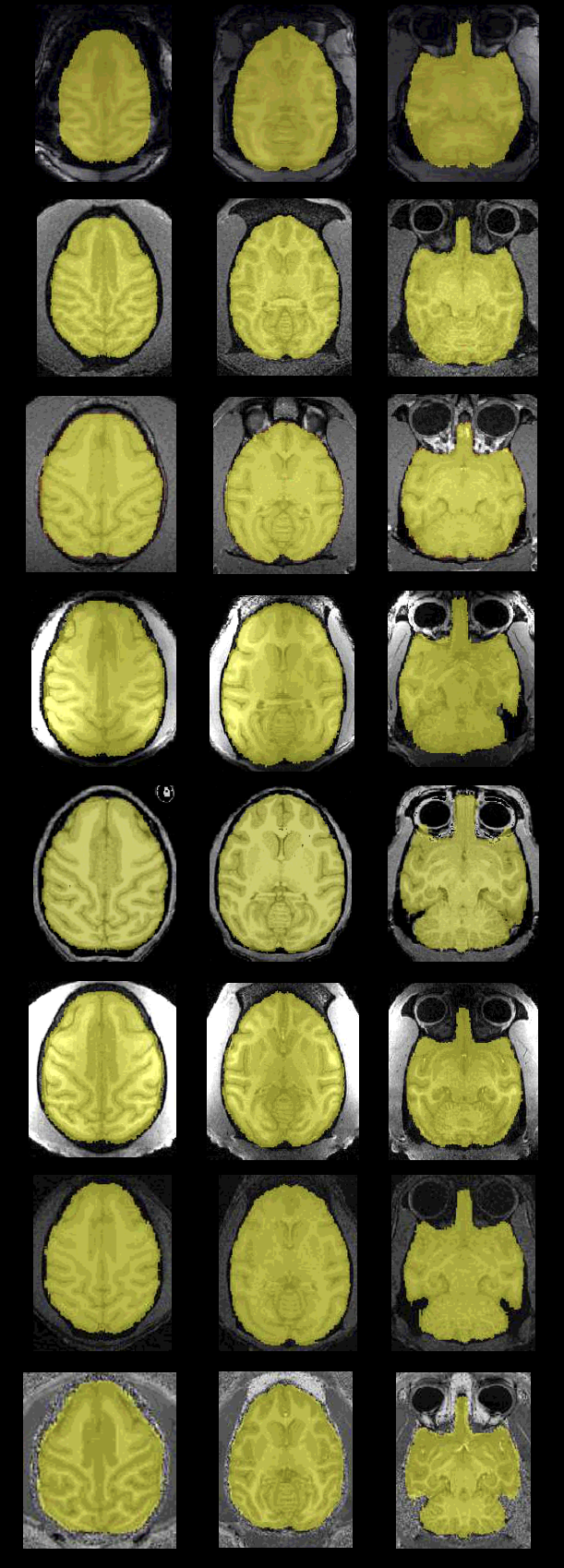

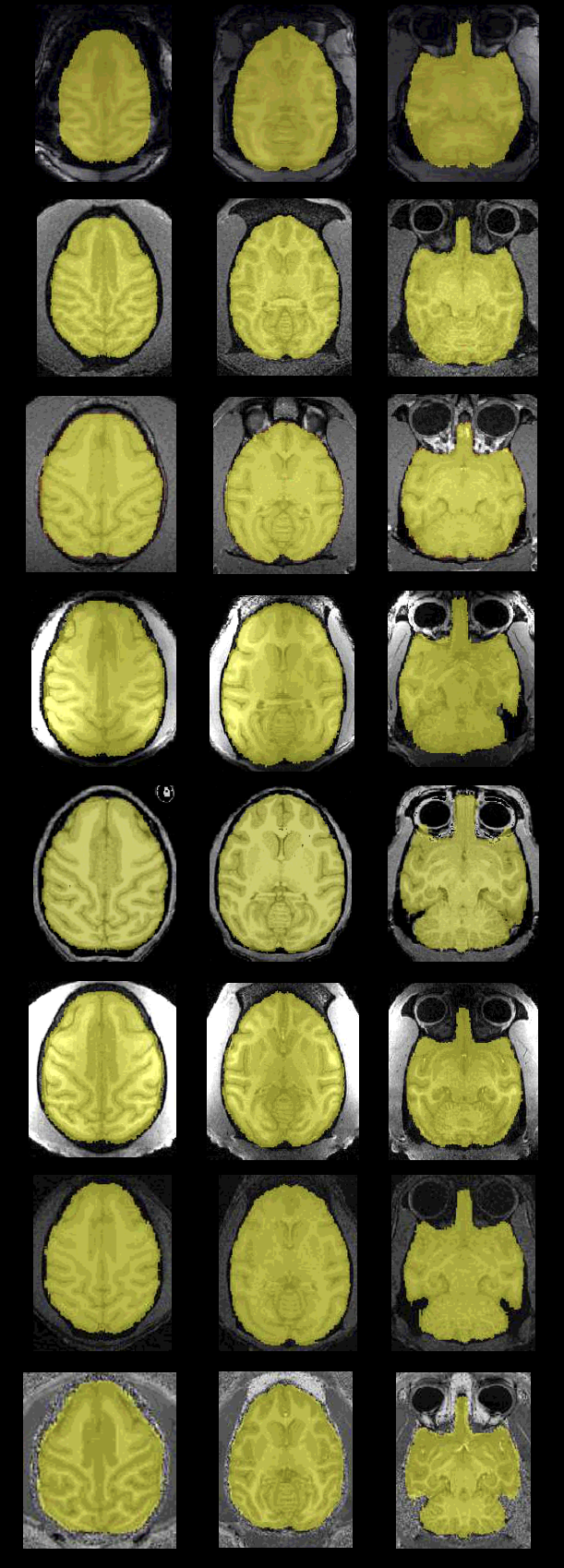

**Supplementary Figure 3- Representative Brainmasks produced by AutoMacq.**

Masks for one subject from each site are illustrated. A: Newcastle, B: DPZ, C: NIDA, D: Oxford, E: MSP, F: MSS, G: SBRI, H: UWO.

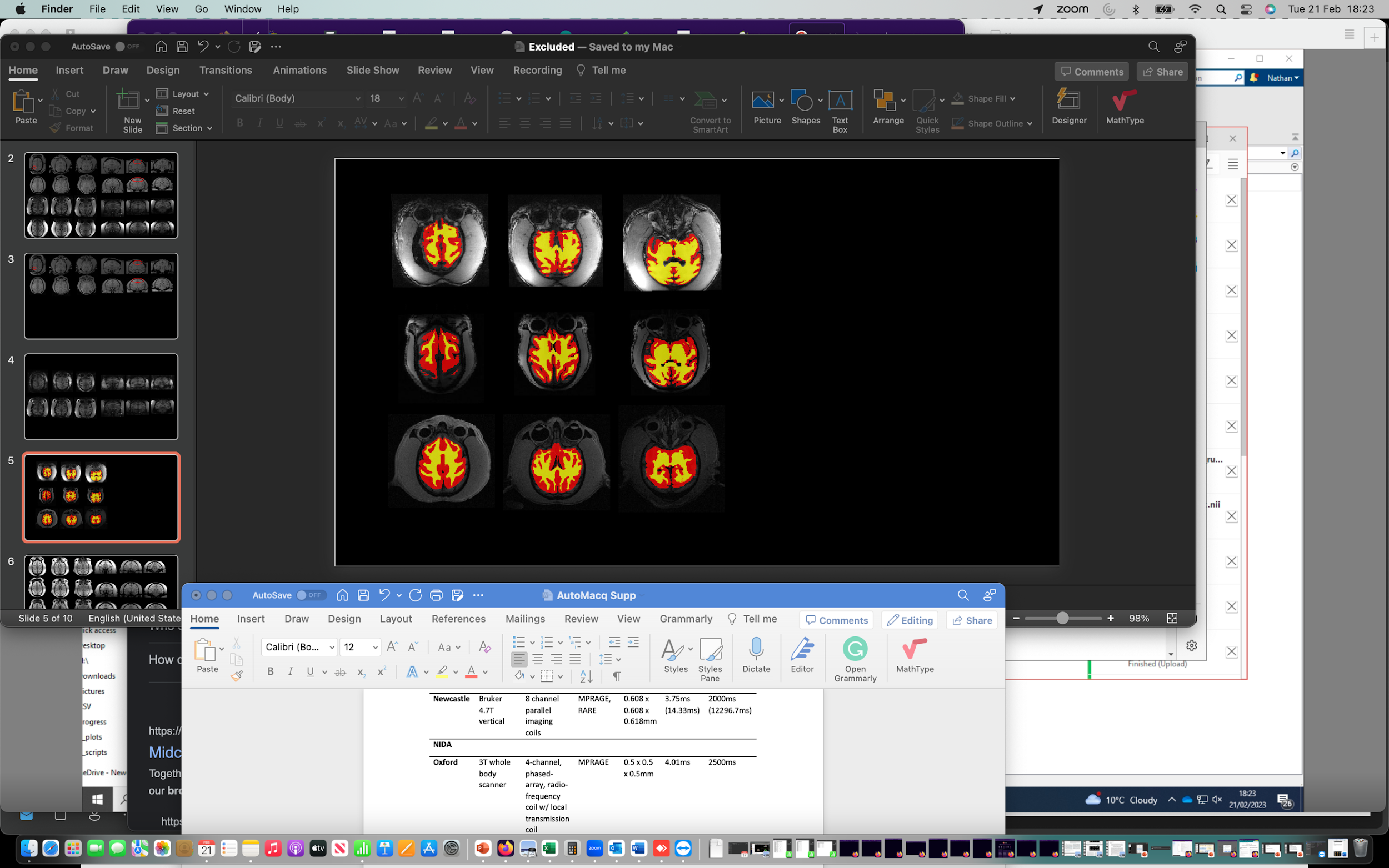

**Supplementary Figure 4- Problematic tissue volumes produced by AutoMacq**. A: subject from the Newcastle dataset, **B:** subject from the Oxford dataset, C: subject is from the DPZ dataset.

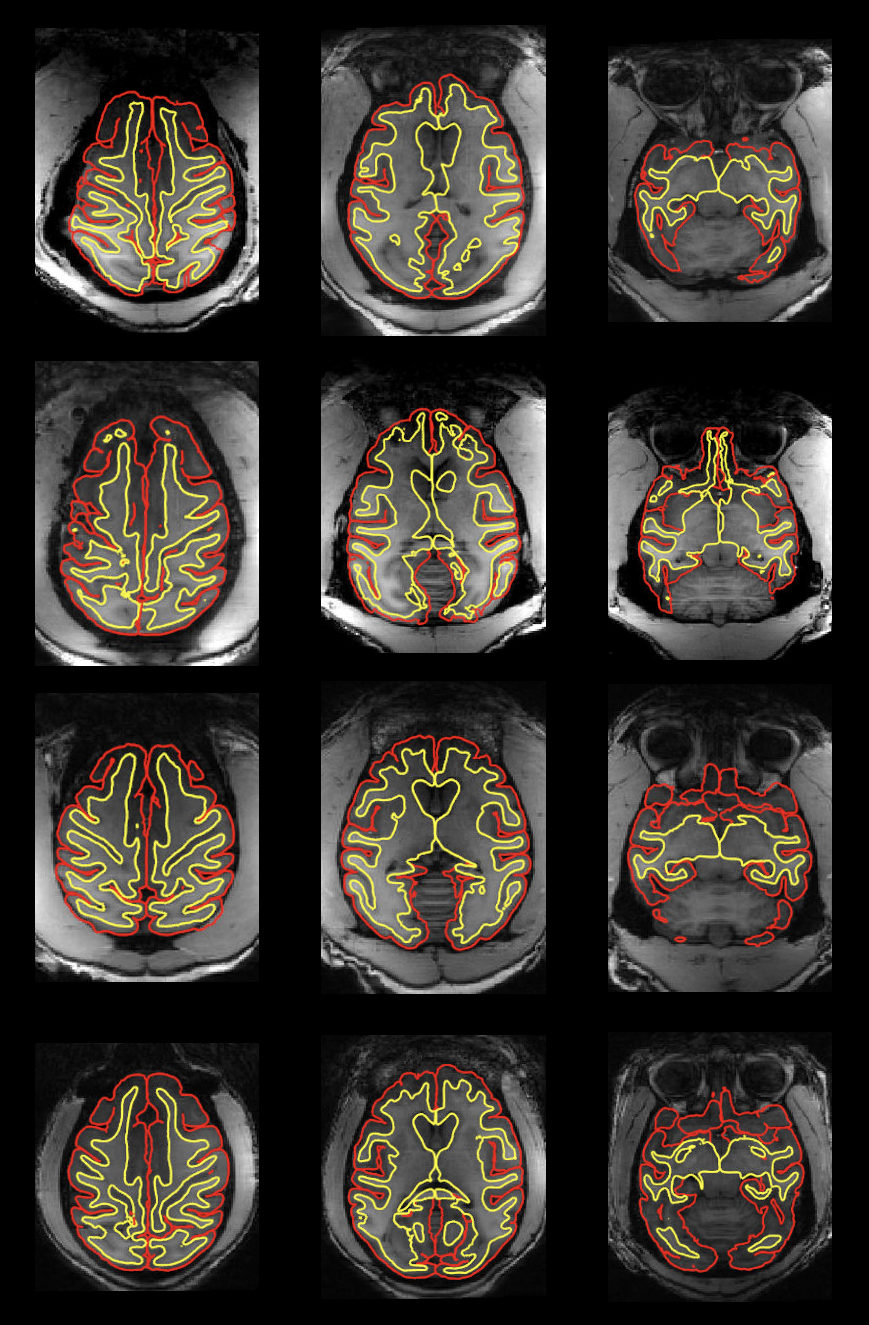

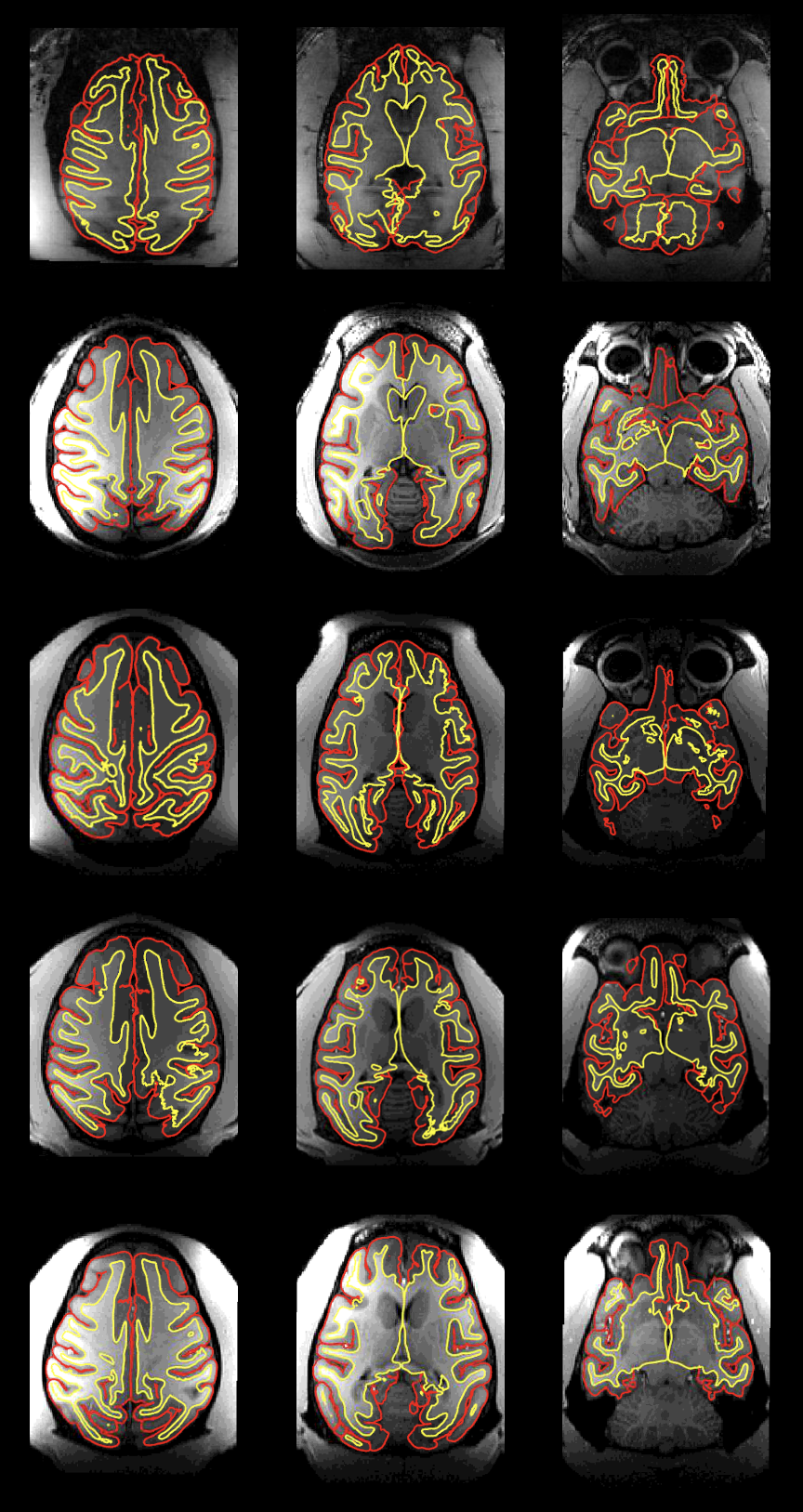

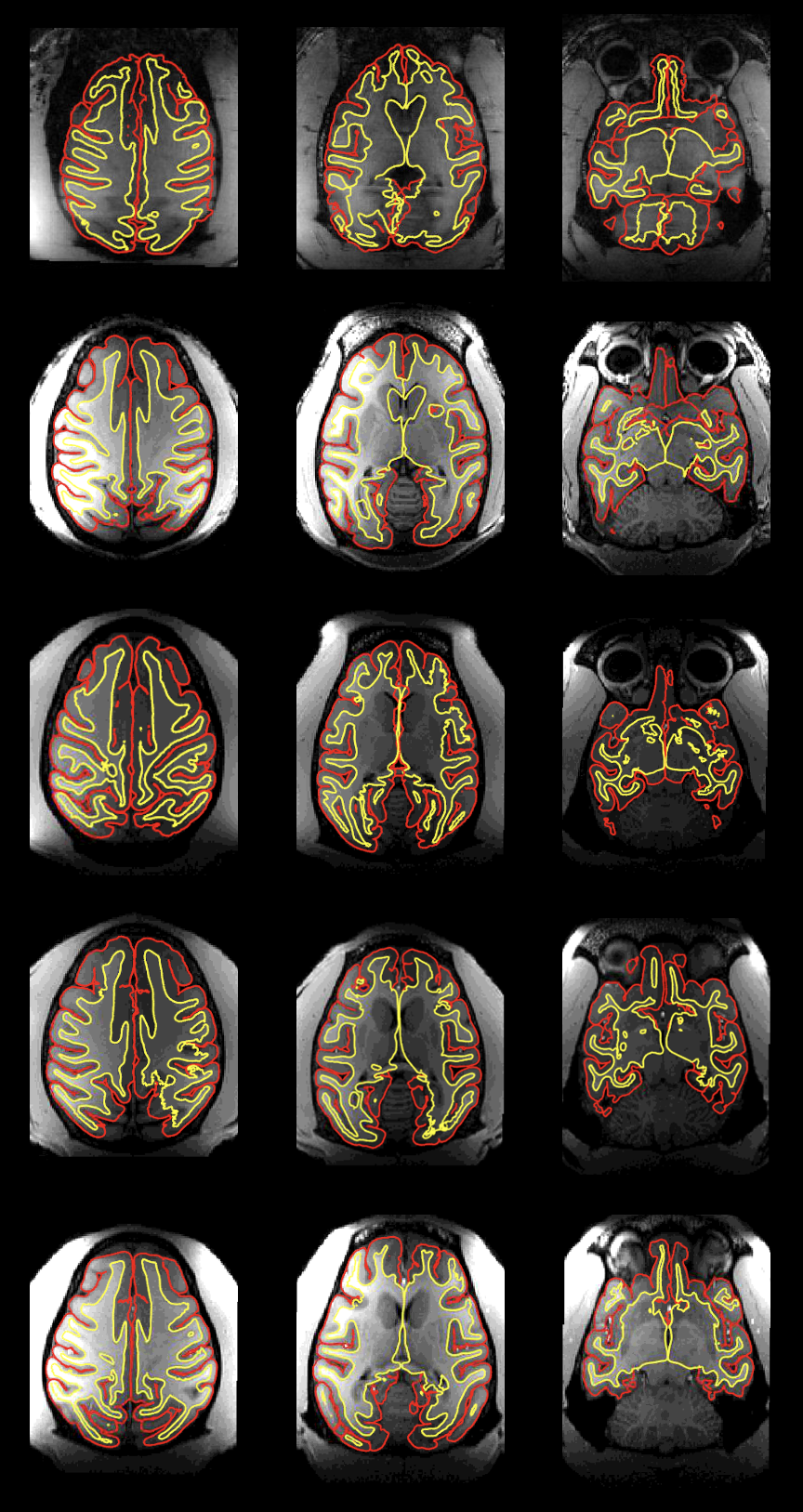

**A4**

**A3**

**A2**

**A1**

**B3**

**B2**

**B1**

**A5**

**B4**

**Supplementary Figure 5-** **Subjects with problematic surfaces produced by AutoMacq.** 5 subjects are from the Newcastle dataset (A) and 4 subjects are from the Oxford dataset (B).
